# A Conserved Interface Between COG4 and Syntaxin 5 Is Essential for Intra-Golgi Membrane Trafficking

**DOI:** 10.64898/2026.11.19.767949

**Authors:** Shegufta Tasneem Neetole, Irina Pokrovskaya, Brian Shank, Walter S. Aragon-Ramirez, Vladimir Lupashin

## Abstract

The conserved oligomeric Golgi (COG) complex tethers retrograde intra-Golgi transport vesicles, but how it engages the SNARE machinery to support vesicle fusion remains unclear. Here, we identify a functional interface between the COG subunit COG4 and the Qa-SNARE Syntaxin 5 (STX5). The C-terminal region of COG4 coprecipitated with STX5, and AlphaFold3 modeling placed the COG4 MUN-C-like subdomain in contact with the STX5 Ha–Hb loop and the linker connecting its Habc domain to the SNARE motif. Replacing the Ha–Hb loop with a flexible Gly/Ser sequence preserved STX5 localization and its interaction with SCFD1 but impaired rescue of GPP130 abundance and protein glycosylation in STX5-depleted cells. Combining the loop replacement with linker substitutions reduced recovery of COG3 and the SNARE partner GS15 with STX5 and led to vesicle accumulation around disorganized Golgi cisternae. On the COG4 side of the interface with Qa-SNARE, alanine substitution of the conserved residues D452 and D453 reduced coprecipitation with STX5. Replacing these residues with either alanine or lysine preserved COG4 localization and COG8 recruitment to the Golgi but failed to restore normal glycosylation, GS15 localization, or altered Golgi morphology in COG4-knockout cells. These findings support a model in which the COG4–STX5 interface couples vesicle tethering to productive SNARE assembly, thereby controlling vesicular retrograde trafficking and Golgi organization.

## Introduction

Eukaryotic cells are compartmentalized, making intracellular vesicular trafficking essential for delivering proteins and lipids to their precise locations. The Golgi apparatus plays a pivotal role in membrane trafficking. After synthesis, up to one-third of proteins enter the secretory pathway and translocate into the ER, from which the majority of secretory proteins are transported through the Golgi (Jackson, 2009), (J. B. Blackburn, D’Souza, & Lupashin, 2019).

Within the Golgi, cargo proteins and lipids are transported and recycled in vesicles with the help of cargo adaptors, coat proteins, small GTPases, soluble N-ethylmaleimide-sensitive factor attachment protein receptors (SNAREs), and vesicular tethers. We focused on SNAREs and tethers because their coordinated activities link vesicle capture to membrane fusion, yet the molecular interactions that couple tethering to SNARE complex assembly remain poorly understood. SNAREs mediate fusion of cargo-containing vesicles with membranes (Y. L. Zhang & Hughson, 2021). They are mostly integral membrane proteins with a characteristic SNARE motif. SNAREs are classified by a single hydrophilic residue in the middle of the SNARE domain, with glutamine designating Q SNAREs and arginine designating R SNAREs. Fusion is mediated by assembly of three Q SNAREs and one R SNARE into a four-helix bundle, designating Q SNAREs and arginine (STX5) as a major bundle; Q SNAREs and arginine (R SNAREs). At the Golgi, the Qa-SNARE syntaxin 5 (STX5) supports vesicular trafficking by participating in several distinct SNARE complexes (D’Souza, Pokrovskaya, & Lupashin, 2023).

CATCHR (complexes associated with tethering containing helical rods) complexes are an evolutionarily conserved family of multisubunit membrane vesicle tethers composed of three to eight distinct subunits. The CATCHR family comprises five complexes. These complexes, which include the yeast Dsl1 (dependent on Sly1-20) complex and its mammalian counterpart NRZ (NBAS–RINT1–ZW10), the conserved oligomeric Golgi (COG) complex, the Golgi-associated retrograde protein (GARP) complex, the endosome-associated recycling protein (EARP) complex, and exocyst, tether transport vesicles at distinct steps in the secretory and endocytic pathways (Bröcker, Engelbrecht-Vandré, & Ungermann, 2010), (Croteau, Furgason, Devos, & Munson, 2009; Ha et al., 2016; Mei et al., 2018; Whyte & Munro, 2002), (Sohda et al., 2010; Sohda et al., 2007; Ungar et al., 2002; Willett, Ungar, & Lupashin, 2013; Y. L. Zhang & Hughson, 2021). The majority of CATCHR subunits contain elongated α-helical bundles (Croteau et al., 2009) and N-terminal coiled-coil regions (Whyte & Munro, 2002). Structural studies of exocyst and the COG complex showed that interactions among these N-terminal regions help assemble subunits into subcomplexes (Whyte & Munro, 2002), (Ha et al., 2016), (Mei et al., 2018). The COG complex is essential for tethering and fusion of intra-Golgi recycling vesicles (Willett et al., 2013). This octameric complex consists of two subcomplexes: lobe A (COG1–COG4), which associates with the Golgi membrane, and lobe B (COG5–COG8) (Ungar et al., 2002), which interacts with vesicular membranes (D’Souza, Taher, & Lupashin, 2020), (Willett et al., 2016). The COG complex coordinates vesicular trafficking via interactions with other critical components, including SNARE proteins, Sec1/Munc18 (SM) proteins, and coiled-coil tethers like Golgin84/GOLGA5 (Sohda et al., 2010), and p115/USO1 (Sohda et al., 2007).

During vesicle trafficking, different CATCHRs interact with pathway-specific SNAREs that regulate vesicle capture, docking, and fusion with the target membrane. Tip20p and Sec39p subunits of the Dsl1 complex bind the N-terminal domains of the Qb SNARE Sec20p and the Qc SNARE Use1p, respectively, anchoring the tether while leaving the SNARE motif available for assembly (Travis et al., 2020). Similarly, NRZ subunits interact with the ER SNAREs STX18, BNIP1, p31, and Sec22b (Tagaya, Arasaki, Inoue, & Kimura, 2014) (Aoki et al., 2009). The N-terminal region of the GARP subunit Vps51p interacts with the Habc domain of the Qc SNARE Tlg1p, paralleling the interaction of human VPS51 with the SNARE STX6 (TerBush, Maurice, Roth, & Novick, 1996). Functional studies have extended these observations beyond mere physical association. In a reconstituted yeast system, purified exocyst, a post-Golgi CATCHR complex, mediates assembly of both binary Q-SNARE and ternary trans-SNARE complexes to induce membrane fusion (Lee et al., 2025). In mammalian cells, GARP depletion reduced assembly of the specific SNARE complex, STX16-STX6-Vti1a-VAMP4, while biochemical studies implicated COG in promoting productive Golgi-SNARE pairing and maintaining the stability of STX5-containing complexes (Pérez-Victoria & Bonifacino, 2009), (Shestakova, Suvorova, Pavliv, Khaidakova, & Lupashin, 2007), (Laufman, Kedan, Hong, & Lev, 2009). Although different CATCHR complexes use similar modes of SNARE recognition, how individual tether-SNARE interactions mediate SNARE activation, assembly, or stability remains an open question.

Previous studies implicated the N-terminal region of COG4 in binding the Sec1/Munc18 (SM) protein SCFD1/Sly1 and the Qa-SNARE STX5, with SCFD1 binding mapped to residues 1–84 and a separate STX5-binding site proposed within residues 84–153 (Shestakova et al., 2007; Laufman et al., 2009). Structural studies subsequently placed the COG4 N-terminus within the assembly core formed by COG1–COG4 (Shu, Jin, Rothman, & Zhang, 2020), raising the question of how this region accommodates STX5 binding in the assembled complex (Shu et al., 2020; Yang et al., 2015).

In this work, we utilize AlphaFold structural modeling to predict a novel protein-protein interface between the COG4 MUN-C-like subdomain and two regulatory regions of STX5, the Ha–Hb loop and the linker connecting the Habc and SNARE domains. Functional analyses in HEK293 and RPE1 cells showed that replacing STX5 Ha–Hb loop residues 66–74 with a flexible Gly/Ser sequence or substituting the evolutionarily conserved COG4 residues D452 and D453, impaired complementation of STX5 depletion or COG4 knockout (KO), respectively. These mutations failed to restore normal Golgi trafficking and glycosylation despite preserving Golgi localization and interactions with other tested protein partners. Together, these findings support a functional interface between COG4 and STX5 that is required for intra-Golgi membrane trafficking and Golgi homeostasis.

## Results

### The C-terminal region of COG4 mediates interaction with STX5

Previous studies implicated the N-terminal region of COG4 (residues 1–222) in STX5 binding (Shestakova et al., 2007), with the proposed binding site subsequently narrowed to residues 84–153 (Laufman et al., 2009). To characterize this interaction, we used an AlphaFold2-predicted structure to identify solvent-exposed residues within this region for mutagenesis (**Supp. Fig. 1A**). We generated four full-length, myc-tagged COG4 mutants containing alanine substitutions at the indicated positions: 3A, D116A K119A D130A; 4A, E106A K111A R113A R121A; 7A-1, D90A K92A E106A K111A R113A D116A K119A; 7A-2, R121A Y123A R128A D130A D131A D134A K136A (**Supp. Fig. 1B**). These substitutions primarily targeted charged surface residues. The COG4 constructs were co-expressed with GFP–STX5 in HEK293 cells, and their association with STX5 was assessed by GFP pull-down. All four mutants retained association with GFP–STX5. Recovery of the 3A, 4A, and 7A-1 mutants was greater than that of WT COG4, although the measurements were variable (**Supp. Fig. 1C, D**). Thus, these substitutions did not reduce COG4 association with STX5 under the overexpression conditions tested. Structural analysis of the yeast COG complex places the COG4 N-terminal region within an assembly formed by the N-termini of COG1, COG2, COG3, and COG4 that stabilizes lobe A (Laufman et al., 2009). This arrangement raises questions about the accessibility of the proposed N-terminal STX5-binding site within the assembled complex and prompted us to investigate additional STX5-binding regions in COG4.

**Figure 1:**
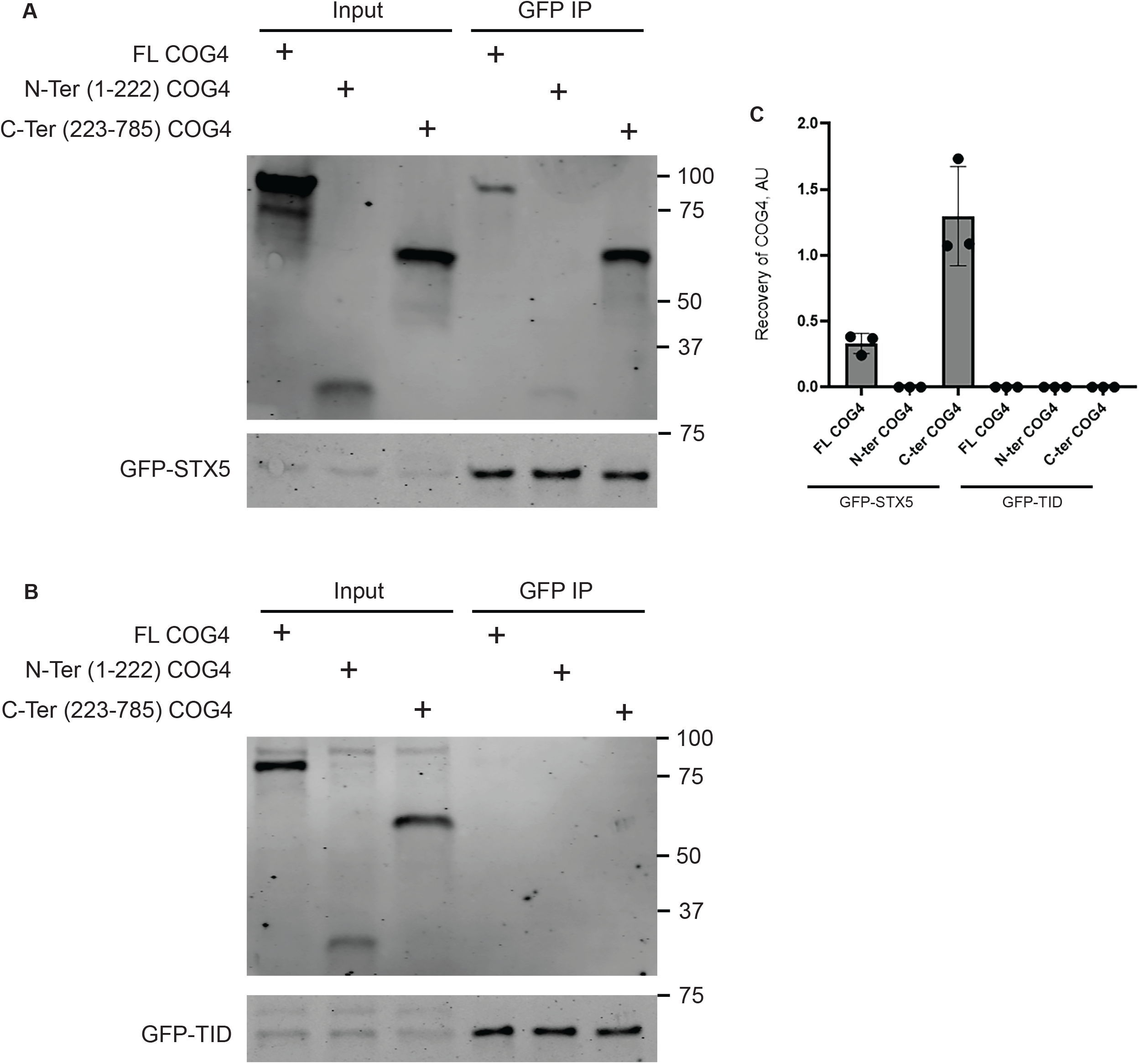
The C-terminal region of COG4 mediates association with STX5. WT HEK293T cells expressing GFP–STX5 (**A**) or GFP–TurboID (GFP–TID; negative control) (**B**) were transiently transfected with plasmids encoding full-length (FL) COG4, the N-terminal fragment (residues 1–222), or the C-terminal fragment (residues 223–785), each bearing a 3×myc tag. Following overnight expression, GFP-STX5 was isolated from cell lysates using GFP-binding protein beads. Input lysates and bound fractions (GFP IP) were analyzed by SDS–PAGE and immunoblotting with anti-myc and anti-GFP antibodies to detect COG4 constructs and GFP-tagged bait proteins, respectively. Molecular masses are indicated in kDa. (**C**) Quantification of COG4 recovery in GFP pulldowns. Anti-myc band intensities were measured using Image Studio and normalized to the corresponding anti-GFP signal in the bound fraction. Values are expressed in arbitrary units (AU) which represent the ratio between the mean signal intensity of COG4 and GFP-STX5. Bars show the mean ± SD from three independent biological replicates, with individual replicates shown as points.

Multiple studies have demonstrated an association between COG4 and STX5 (Laufman et al., 2009; Shestakova et al., 2007). Because the COG4 N-terminal region participates extensively in interactions within the COG complex, we investigated whether its C-terminal region contributes to STX5 binding. We used AlphaFold2 and AlphaFold3 (AF3) (Jumper et al., 2021), (Abramson et al., 2024) to model COG–STX5 interactions using protein sequences from humans, mice, fish, frogs, octopuses, plants, worms, and yeast (data not shown). We also included SCFD1/Sly1 in these analyses because STX5 and its yeast homolog Sed5 are bound and regulated by this SM protein (Laufman et al., 2009), (Duan et al., 2024), (Peng & Gallwitz, 2002), (Dulubova et al., 2003). Analysis of the AF3-predicted structure identified COG4 as the only COG subunit interacting with STX5 through its C-terminal region (data not shown).

Previous comparative sequence and structural analyses identified similarities between several CATCHR subunits, including COG4, and the MUN domain of UNC13/Munc13 (Pei, Ma, Rizo, & Grishin, 2009). To examine this relationship, we compared the AF3-predicted structures of the MUN domain of the human UNC13A (residues 920–1394) and COG4 (residues 134–785) using TM-align (Y. Zhang & Skolnick, 2005). Structural alignment yielded an RMSD of 4.91 Å and a TM-score of 0.671, supporting a common fold (**Supp. Fig. 2C**). Together with the established role of UNC13/Munc13 in Syntaxin 1 regulation and SNARE assembly (Y. Zhang & Skolnick, 2005). This structural similarity provided a rationale for investigating the COG4 MUN-like region as a functional STX5-binding module. To test this hypothesis, we expressed myc-tagged full-length COG4 (FL), its N-terminal fragment (residues 1–222), or its C-terminal fragment (residues 223–785) in HEK293T cells stably expressing GFP–STX5. GFP–STX5 and associated proteins were isolated using GFP-binding protein (GBP) beads. Full-length COG4 was coprecipitated with GFP–STX5, whereas recovery of the N-terminal fragment was weak. The C-terminal fragment showed the greatest recovery, exceeding that of full-length COG4 after normalization to the GFP–STX5 signal (**Fig. 1A, C**). As a specificity control, stably expressed GFP–TurboID did not recover any of the COG4 constructs. (**Fig. 1B**). These results support a specific association between the COG4 C-terminal region and STX5, consistent with structural prediction.

**Figure 2:**
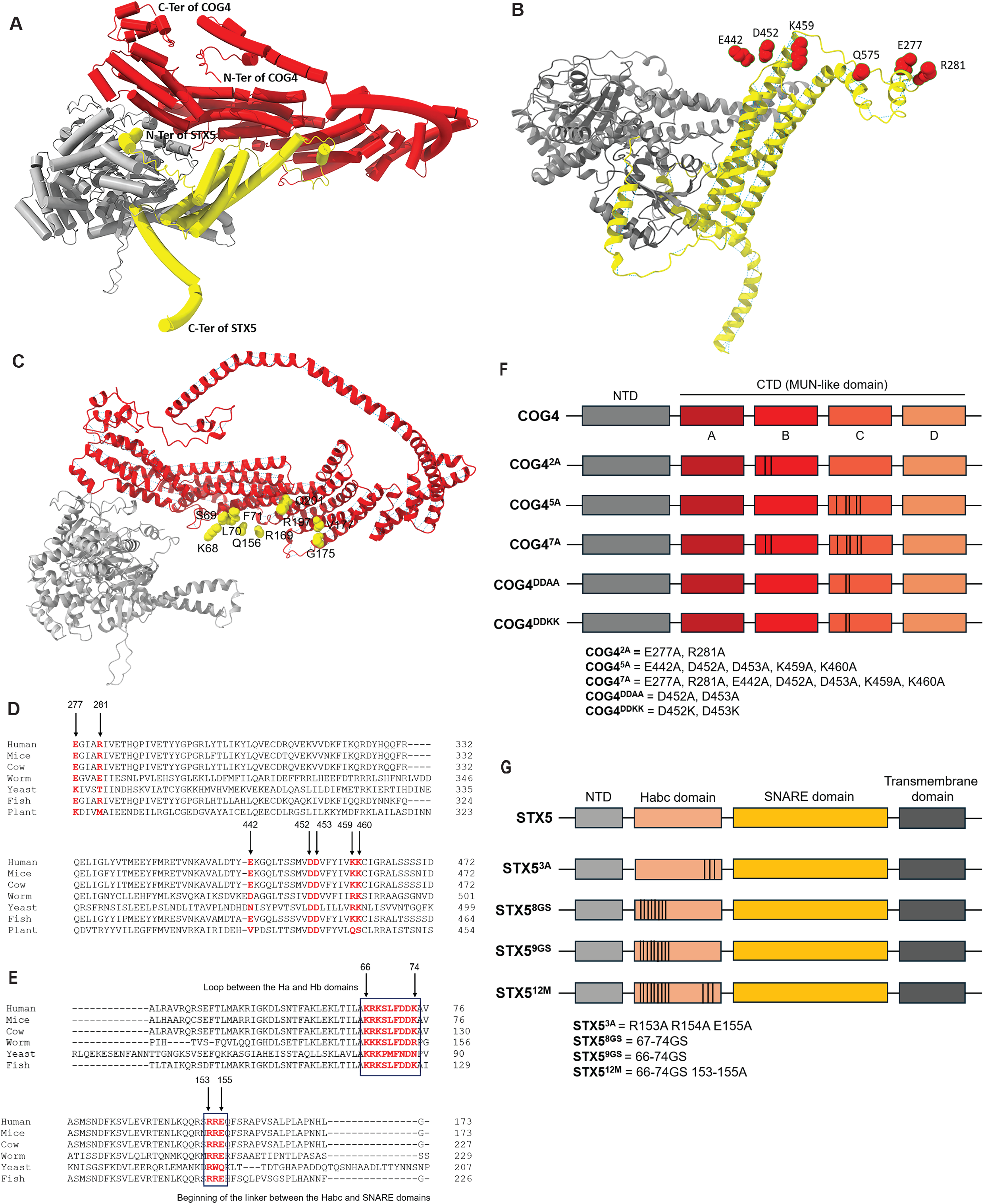
**AlphaFold3 predicts an interface between the COG4 MUN-C-like subdomain and regulatory regions of STX5**. **(A)** Predicted ternary complex of COG4 (red), STX5 (yellow), and SCFD1 (gray). The N- and C-termini of COG4 and STX5 are indicated. (**B**) View highlighting selected COG4 residues examined by mutagenesis, shown as red spheres. STX5 is shown in yellow and SCFD1 in gray. (**C**) Complementary view highlighting selected STX5 residues near the predicted COG4 interface, shown as yellow spheres. COG4 is shown in red and SCFD1 in gray. (**D**) Multiple sequence alignments of COG4 regions containing E277/R281 and E442/D452/D453/K459/K460. Residues selected for mutagenesis are highlighted in red, with human residue numbers indicated above the alignments. (**E**) Multiple sequence alignments of the STX5 Ha–Hb loop and the beginning of the linker connecting the Habc and SNARE domains. Boxes identify the regions targeted by mutagenesis, including residues 66–74 and R153/R154/E155. Numbering corresponds to human STX5 isoform 2. (**F**) Domain organization of COG4 and the mutants used in this study. The N-terminal domain (NTD) and C- terminal MUN-like subdomains A–D are indicated. COG4^2A^ contains E277A/R281A; COG4^5A^ contains E442A/D452A/D453A/K459A/K460A; COG4^7A^ combines the 2A and 5A substitutions; COG4ᴰᴰᴬᴬ contains D452A/D453A; and COG4ᴰᴰᴷᴷ contains D452K/D453K. (**G**) Domain organization of STX5 and the mutants used in this study. STX5^3A^ contains R153A/R154A/E155A; STX5^8GS^ and STX5^9GS^ replace residues 67–74 and 66–74, respectively, with Gly/Ser sequences; and STX5^12M^ combines the 9GS replacement with R153A/R154A/E155A. Vertical lines in panels **F** and **G** indicate the approximate mutation positions.

### AlphaFold3 predicts an interface between the COG4 MUN-C subdomain and STX5

We used AF3 to model a ternary complex of COG4, STX5, and SCFD1 (**Fig. 2A**). Using the AF3-predicted structure, we identified candidate COG4-STX5 interacting residues. From the COG4 side, E277, R281, E442, D452, K459, and Q575 were identified as possibly being involved in this interaction (**Fig. 2B**). Predicted STX5 interface residues included R67, K68, S69, L70, and F71 in the Ha–Hb loop, Q156, R169, G175, V177, and R197 in the Habc–SNARE linker, and Q201 near the beginning of the SNARE motif (**Fig. 2C**). To further validate the biological relevance of the predicted interaction interface, we conducted multiple sequence alignments of COG4 and STX5 (isoform 2) (**Fig. 2D, E**), which shows strong evolutionary conservation of the proposed residues. A more detailed representation of the proposed interface is shown in **Supplementary Fig. 3**.

The C-terminal region of COG4 adopts a predicted MUN-like fold comprising four α-helical subdomains, A–D, resembling the MUN domain of Munc13-1, which promotes neuronal SNARE assembly (Xu et al., 2017). In the COG4–STX5 model, the MUN-C-like subdomain (residues 427-624) contacts the Ha–Hb loop of the STX5 Habc domain and the linker connecting Habc to the SNARE motif. Guided by these predictions and sequence conservation, we generated five COG4 mutants and four STX5 mutants targeting the proposed interface. Their domain organization and mutation sites are summarized in **Fig. 2F and G**.

### Mutations in the predicted COG4-binding surface of STX5 impair glycosylation

To assess the functional importance of the predicted COG4-binding surface of STX5, we established a complementation system in which endogenous STX5 was depleted in cells stably expressing GFP-tagged STX5 variants. This strategy allowed us to examine mutant function while accommodating the requirement for STX5 in cell growth (Dascher, Matteson, & Balch, 1994), (D’Souza et al., 2023). Endogenous STX5 was depleted using a previously published siRNA targeting the sequence TAGCCTCAACAAACAAATT (D’Souza et al., 2023). The GFP–STX5 constructs were designed to be siRNA resistant by introducing five synonymous nucleotide substitutions within the target sequence. Guided by the AF3-predicted interface (**Fig. 2C**), we generated four mutants of human STX5 isoform 2, which preferentially functions in the Golgi (Hui et al., 1997), (Linders et al., 2021). STX5^3A^ contains R153A/R154A/E155A alanine substitutions at the beginning of the Habc–SNARE linker. STX5^8GS^ replaces Ha–Hb loop residues 67–74 with an eight-residue flexible Gly/Ser sequence, whereas STX5^9GS^ replaces the entire loop, residues 66–74, with a nine-residue Gly/Ser sequence. STX5^12M^ combines the 9GS replacement with R153A/R154A/E155A. Each construct carried an N-terminal GFP tag and was stably expressed in HEK293 cells under the control of the STX5 promoter region (D’Souza et al., 2023) before depletion of endogenous STX5. Cells stably expressing siRNA-resistant wild-type (WT) GFP–STX5 served as the positive complementation control. To assess whether the mutations affected STX5 localization, we examined the GFP– STX5 constructs in the resulting cell lines. All constructs localized to the Golgi, as indicated by colocalization with GM130/GOLGA2, suggesting that the substitutions preserved Golgi targeting (**Supp. Fig. 4**). We also compared GFP–STX5^WT^ and GFP–STX5^12M^, the most extensively mutated variant, for association with coexpressed HA-tagged SCFD1. GFP pulldown recovered similar amounts of HA–SCFD1 with both constructs, supporting preservation of the SCFD1-binding conformation in the STX5^12M^ mutant (**Supp. Fig. 5**).

**Figure 3.**
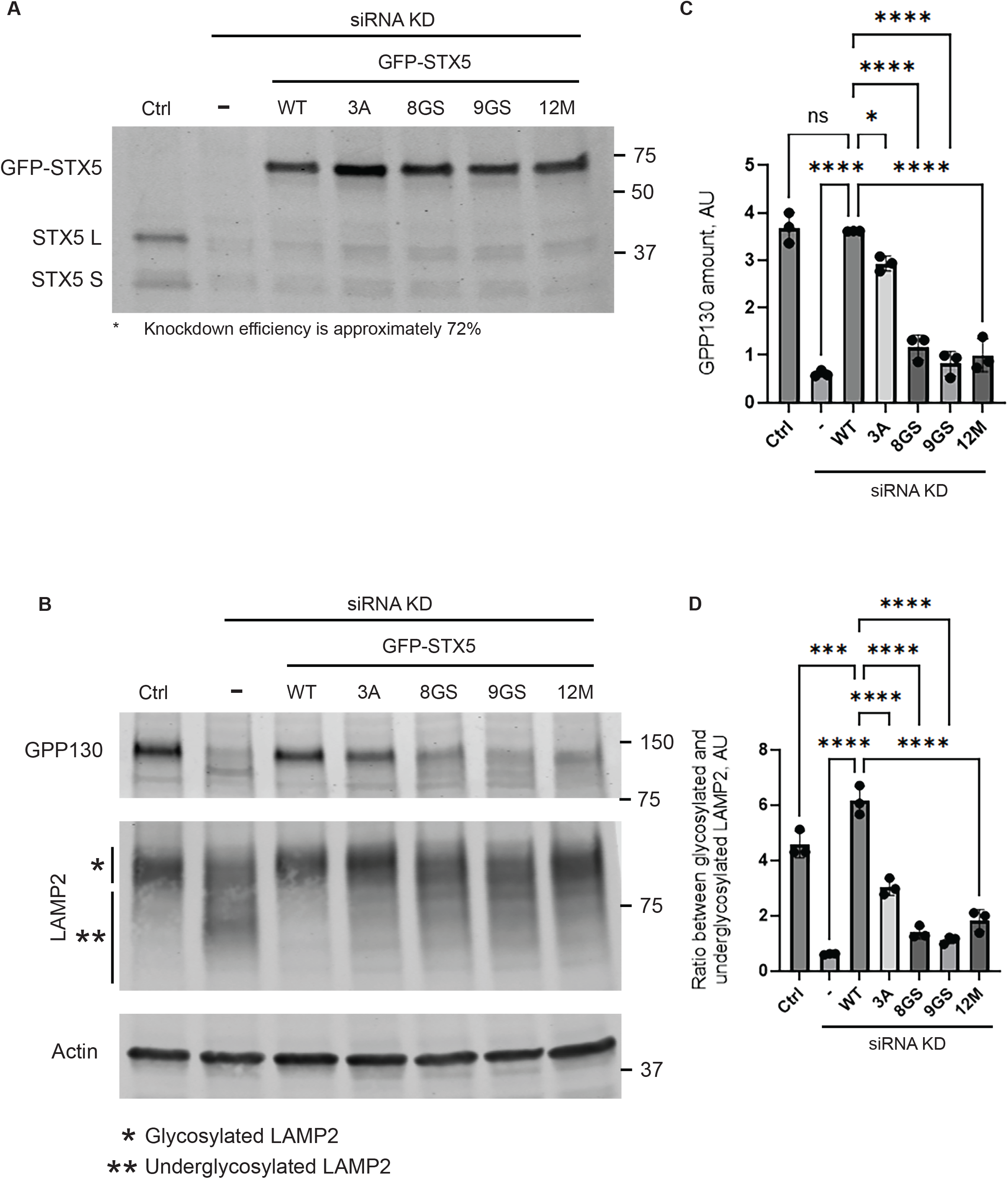
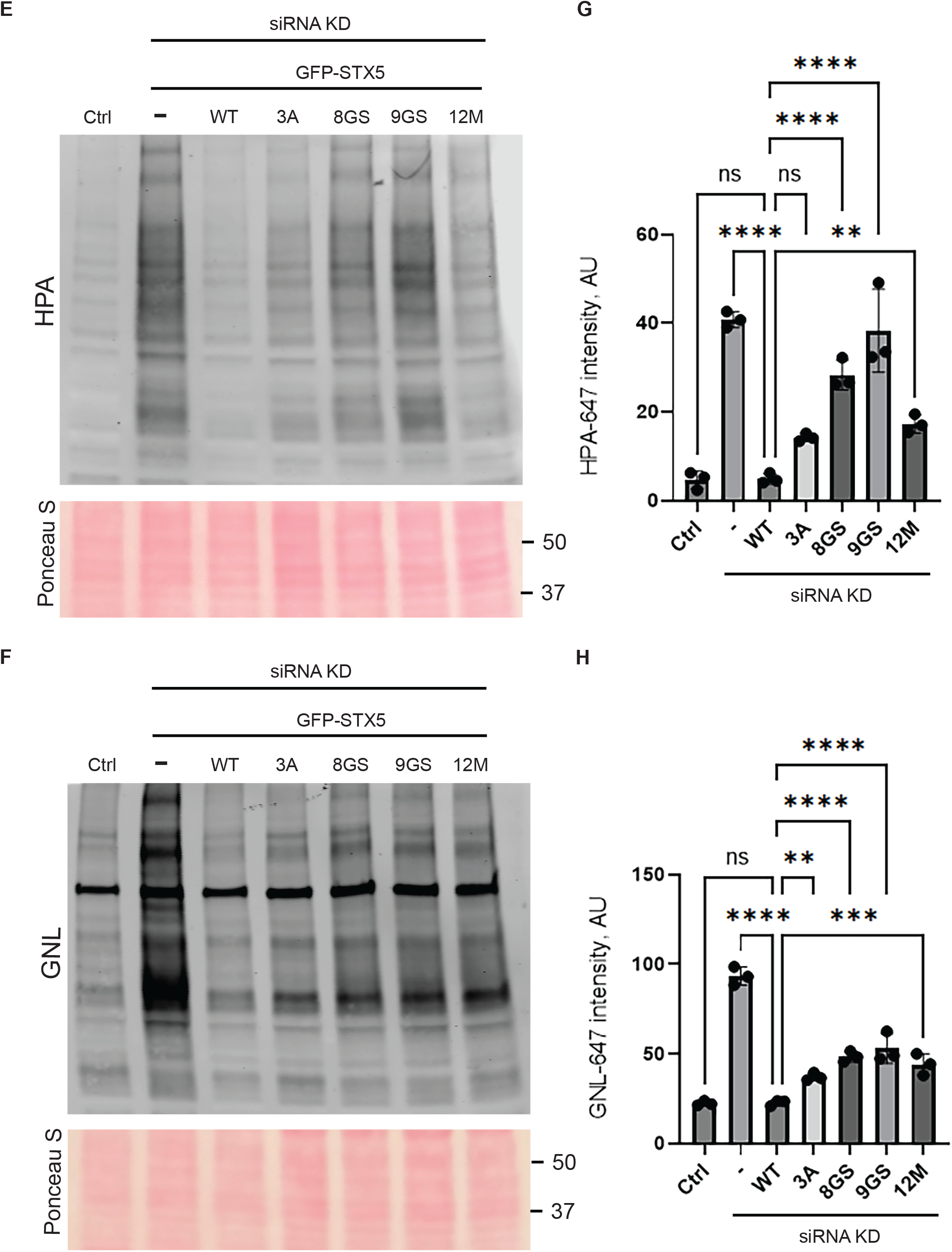
Mutations within the predicted COG4-binding surface of STX5 impair Golgi glycosylation and GPP130 maintenance expression. Experiments were performed in WT HEK293T cells or HEK293T cells stably expressing siRNA-resistant GFP–STX5 isoform 2, either wild-type (WT) or the indicated mutants. Ctrl, wild-type HEK293 cells without STX5 knockdown; −, WT HEK293 cells depleted of STX5 without a complementation construct. Mutant definitions are provided in Figure 2. (**A**) Immunoblot analysis with an anti-STX5 antibody showing depletion of endogenous long (STX5 L) and short (STX5 S) isoforms and expression of GFP–STX5 constructs. (**B**) Immunoblot analysis of GPP130/GOLIM4 and LAMP2 under the same experimental conditions. Actin serves as a loading control. Increased electrophoretic mobility of LAMP2 is consistent with underglycosylation. (**C**) Quantification of GPP130 abundance, normalized to actin. **(D)** Quantification of the ratio of glycosylated to underglycosylated LAMP2. (**E–H**) HPA and GNL lectin blots and quantification of lectin binding, assessing alterations in O- and N-linked glycosylation, respectively. Membranes were probed with fluorescently labeled HPA-647 or GNL-647 and analyzed by fluorescence scanning. Corresponding Ponceau S–stained membranes are shown beneath the lectin blots to verify comparable total protein loading. Band and lectin signal intensities were measured using LI-COR Image Studio version 5.5, and graphs were generated using GraphPad Prism 10. Molecular masses are indicated in kDa. Bars show mean ± SD from three independent biological replicates, with individual replicates shown as points. Statistical comparisons are indicated by brackets. ns, not significant; *P < 0.05; ***P < 0.001; ****P < 0.0001.

**Figure 4.**
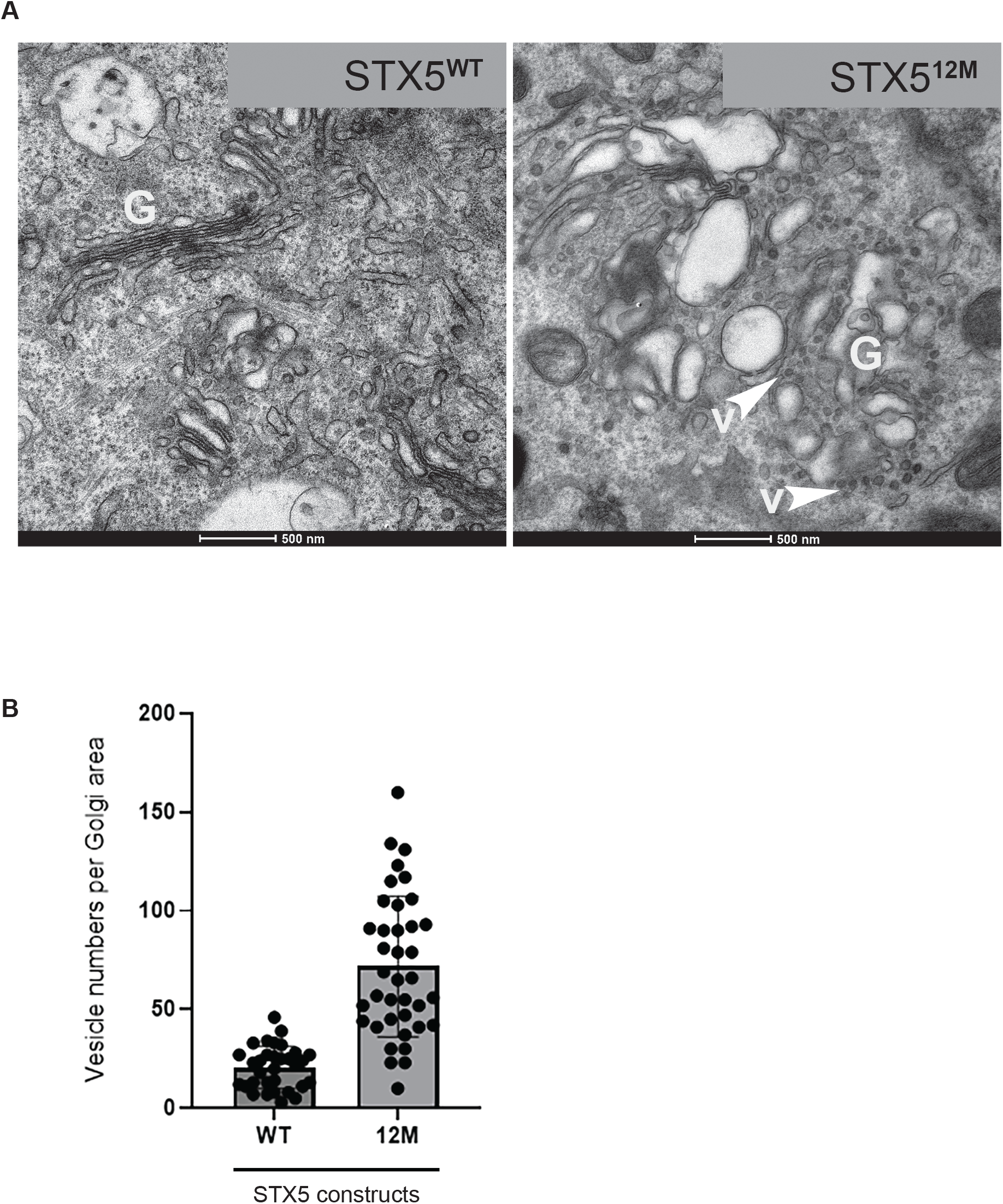
Mutations within the predicted COG4-binding surface of STX5 cause accumulation of Golgi-associated vesicles and Golgi disorganization. (A) Representative transmission electron micrographs (TEM) of HEK293T cells stably expressing siRNA-resistant GFP–STX5 isoform 2, either wild-type (WT) or the STX5^12M^ mutant, following siRNA-mediated depletion of endogenous STX5. Stable expression was established before STX5 depletion. Five days after siRNA transfection, cells were fixed, embedded in Araldite 502/Embed 812 resins, and processed for TEM. The STX5^12M^ mutant combines replacement of Ha–Hb loop residues 66–74 with a Gly/Ser sequence and the R153A/R154A/E155A substitutions. Cells expressing the STX5^12M^ mutant show increased vesicle numbers near the Golgi and disorganized Golgi membranes. G, Golgi apparatus; V and arrowheads, vesicles. Scale bars, 500 nm. (B) Quantification of vesicles in images of Golgi regions acquired at 14,500× magnification. Each point represents the vesicle count in one image. Bars show mean ± SD. Graphs were generated using GraphPad Prism 10.

**Figure 5.**
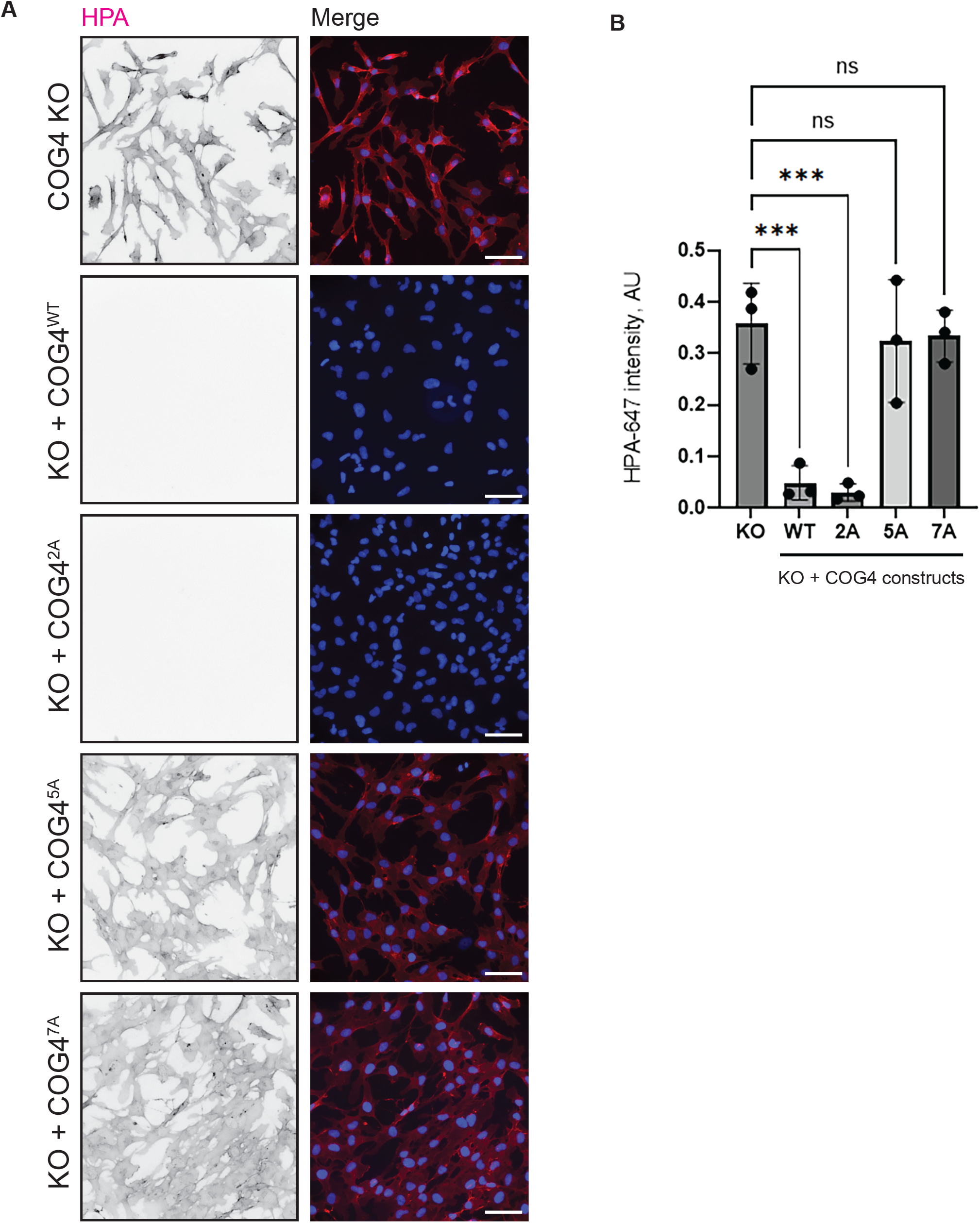
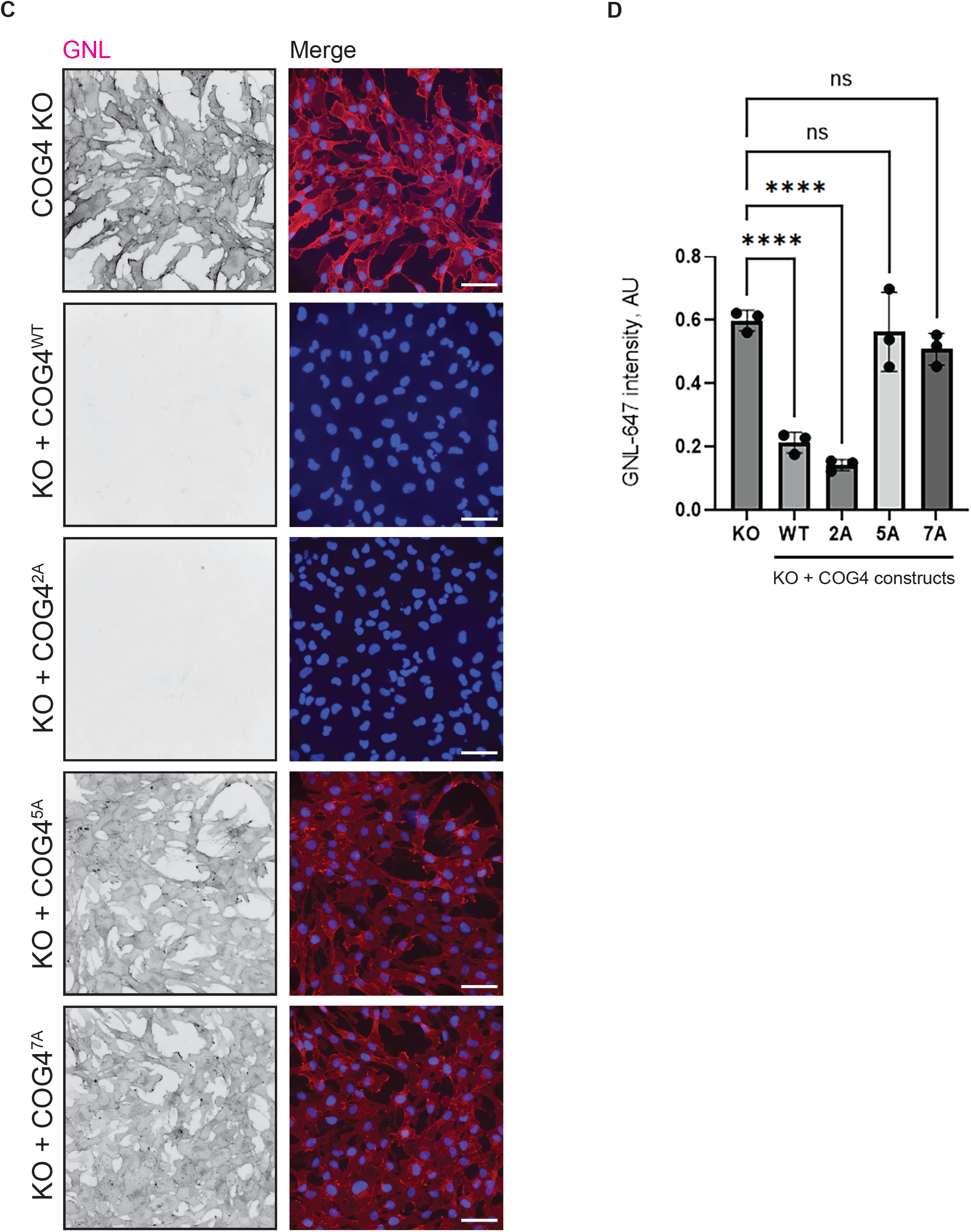
**COG4 interface mutants COG4^5A^ and COG4^7A^ fail to correct glycosylation defects in COG4-knockout cells**. (**A, C**) Representative fluorescence images of RPE1 COG4-knockout (KO) cells and KO cells complemented with myc-tagged wild-type COG4 (WT) or the indicated COG4 mutants (COG4^2A^, COG4^5A^, and COG4^7A^), stained with HPA–647 (**A**) or GNL–647 (**C**) to assess alterations in O- and N-linked glycosylation, respectively. Mutant definitions are provided in Figure 2. Lectin signals are displayed in inverted grayscale in individual channels and in RGB in merged images; nuclei are shown in blue. Scale bars, 20 µm. (**B, D**) Quantification of HPA (**B**) and GNL (**D**) fluorescence. Mean fluorescence intensity was measured for each image using ImageJ and divided by the number of cells in that image. Values are expressed in arbitrary units (AU). Bars show mean ± SD, with individual measurements shown as points. Differences among groups were assessed by one-way ANOVA in GraphPad Prism, yielding overall P values of 0.0002 for HPA and <0.0001 for GNL. Pairwise comparisons are indicated by brackets. ns, not significant; ***P < 0.001; ***P < 0.0001

Following siRNA treatment, both endogenous STX5 isoforms, STX5-L and STX5-S, were depleted, whereas expression of the siRNA-resistant GFP–STX5 constructs was maintained (**Fig. 3A**). Quantification of immunoblotting indicated 72% depletion of endogenous STX5 (**Fig. 3A**). We next assessed GPP130/GOLIM4, a Golgi integral membrane protein whose steady-state abundance is sensitive to COG dysfunction (Oka, Ungar, Hughson, & Krieger, 2004). GPP130 localizes to the cis/medial Golgi and cycles between the Golgi and endosomal compartments (Linstedt, Mehta, Suhan, Reggio, & Hauri, 1997), (Oka et al., 2004). STX5 depletion markedly reduced GPP130 abundance (**Fig. 3B**, lane 2, and **Fig. 3C** for quantification), consistent with STX5’s necessity in recycling this protein to the Golgi. Wild-type GFP–STX5 restored GPP130 abundance to control levels (**Fig. 3B**, lane 3), showing that GFP-tagged STX5 isoform 2 can complement depletion of both endogenous STX5 isoforms. GFP–STX5^3A^ also substantially restored GPP130 abundance (**Fig. 3B**, lane 4), indicating that the R153A/R154A/E155A substitutions largely preserve this function. In contrast, GPP130 abundance remained significantly reduced in cells expressing GFP–STX5^8GS^, GFP–STX5^9GS^, or GFP–STX5^12M^ (**Fig. 3B**, lanes 5–7; **Fig. 3C**). These results identify the STX5 Ha–Hb loop as an important determinant of recycling of Golgi proteins.

COG deficiency impairs glycosylation of proteins traversing the secretory pathway, including the lysosomal membrane protein LAMP2 (J. Blackburn, Pokrovskaya, & Lupashin, 2018). Mature, heavily glycosylated LAMP2 migrates at approximately 120 kDa, whereas underglycosylation increases its electrophoretic mobility, providing a readout of glycosylation defects. STX5-depleted cells without a complementation construct (−) showed a downward shift in LAMP2 migration, whereas untreated control cells (Ctrl) and cells expressing wild-type GFP– STX5 predominantly retained the mature LAMP2 pattern (**Fig. 3B**). Each STX5 mutant showed a downward shift to varying degrees, indicating differences in its ability to restore LAMP2 glycosylation. Quantification of the ratio of glycosylated to underglycosylated LAMP2 is shown in **Fig. 3D**. (Oka et al., 2004) COG-deficient cells exhibit severe O-and N-glycosylation defects caused by impaired recycling of the Golgi glycosylation machinery (Sumya, Aragon-Ramirez, & Lupashin, 2025). We therefore assessed glycosylation as a functional readout of Golgi trafficking using fluorescently labeled Helix pomatia agglutinin (HPA–647) and Galanthus nivalis lectin (GNL–647). HPA recognizes terminal α-N-acetylgalactosamine, specifically the Tn epitope on O-glycoproteins, whereas GNL recognizes terminal mannose residues on N-linked glycans (Sumya, Pokrovskaya, & Lupashin, 2021). STX5 depletion significantly increased binding of both lectins to cellular glycoproteins, and expression of wild-type GFP–STX5 restored lectin binding to control levels (**Fig. 3E–H**). GFP–STX5^3A^ partially complemented the defect, indicating that the R153A/R154A/E155A substitutions retain substantial activity in maintaining Golgi glycosylation. In contrast, cells expressing GFP–STX5^8GS^, GFP–STX5^9GS^, or GFP–STX5^12M^ retained significant O- and N-glycosylation defects (**Fig. 3E–H**). Together, these results further support the requirement of the STX5 Ha–Hb loop in the Golgi trafficking processes that sustain normal glycosylation.

N-ethylmaleimide (NEM) treatment inhibits STX5-containing SNARE complex disassembly, increases STX5 association with COG, and reduces its association with SCFD1 (D’Souza et al., 2023). To determine whether STX5^12M^ responds similarly, we isolated GFP–STX5 from cells stably expressing either the WT or 12M construct following NEM treatment. Both constructs showed more than 90% reduced association with SCFD1 after treatment (**Supp. Fig. 5**). However, GFP–STX5^12M^ recovered two times less GS15/BET1L and COG3 than WT GFP–STX5 under these conditions. These findings suggest that disruption of the predicted COG4-binding interface compromises STX5 in association with COG and the assembly or stability of GS15-containing intra-Golgi SNARE complexes.

### Disruption of the COG4-STX5 interface causes accumulation of Golgi-associated vesicles

Accumulation of Golgi-derived vesicles followed by Golgi fragmentation is a hallmark of acute COG deficiency (Zolov & Lupashin, 2005). Therefore, disruption of the predicted COG4–STX5 interface should produce a similar phenotype if this interaction is indeed needed for vesicular tethering. To assess this, endogenous STX5 was depleted in HEK293 cells stably expressing siRNA-resistant GFP–STX5^WT^ or GFP–STX5^12M^, and Golgi ultrastructure was examined by transmission electron microscopy. Cells expressing GFP–STX5WT retained well-organized Golgi stacks, supporting the tagged protein’s function (**Fig. 4A**). In contrast, cells expressing GFP–STX5^12M^ showed swollen, disorganized Golgi cisternae with marked accumulation of vesicular profiles (**Fig. 4A, B**). Many vesicles were closely opposed to Golgi membranes, consistent with impaired vesicle fusion.

Together, the complementation experiments show that mutations in the predicted COG4-binding interface compromise STX5 function despite preserving its Golgi localization and SCFD1 association. Ha–Hb loop mutants failed to maintain normal GPP130 abundance and Golgi glycosylation. The combined STX5^12M^ mutant also showed reduced recovery of COG3 and GS15 and failed to preserve Golgi ultrastructure. These findings support a role for the predicted COG4–STX5 interface in coupling COG function to productive intra-Golgi SNARE complex assembly and membrane fusion.

### COG4^5A^ and COG4^7A^ fail to restore normal glycosylation in COG4-knockout RPE1 cells

Guided by AF3 predictions and sequence alignments of COG4 homologs, we generated three myc-tagged COG4 mutants. COG4^2A^ carries E277A/R281A alanine substitutions in the MUN-B-like subdomain; COG4^5A^ carries E442A/D452A/D453A/K459A/K460A substitutions in the MUN-C-like subdomain, and COG4^7A^ combines both sets of substitutions (**Fig. 2F**). We generated stable hTERT-RPE1 COG4-KO cell lines expressing wild-type COG4 or one of the three mutants under the control of the COG4 promoter (Sumya et al., 2021). We chose hTERT-RPE1 cells because their growth and Golgi function are more sensitive to COG dysfunction than those of HEK293 cells (!!! INVALID CITATION !!!). Immunofluorescence showed that wild-type COG4 and all three mutants were localized to the Golgi area, colocalizing with GM130 (**Supp. Fig. 6A, B**).

**Figure 6.**
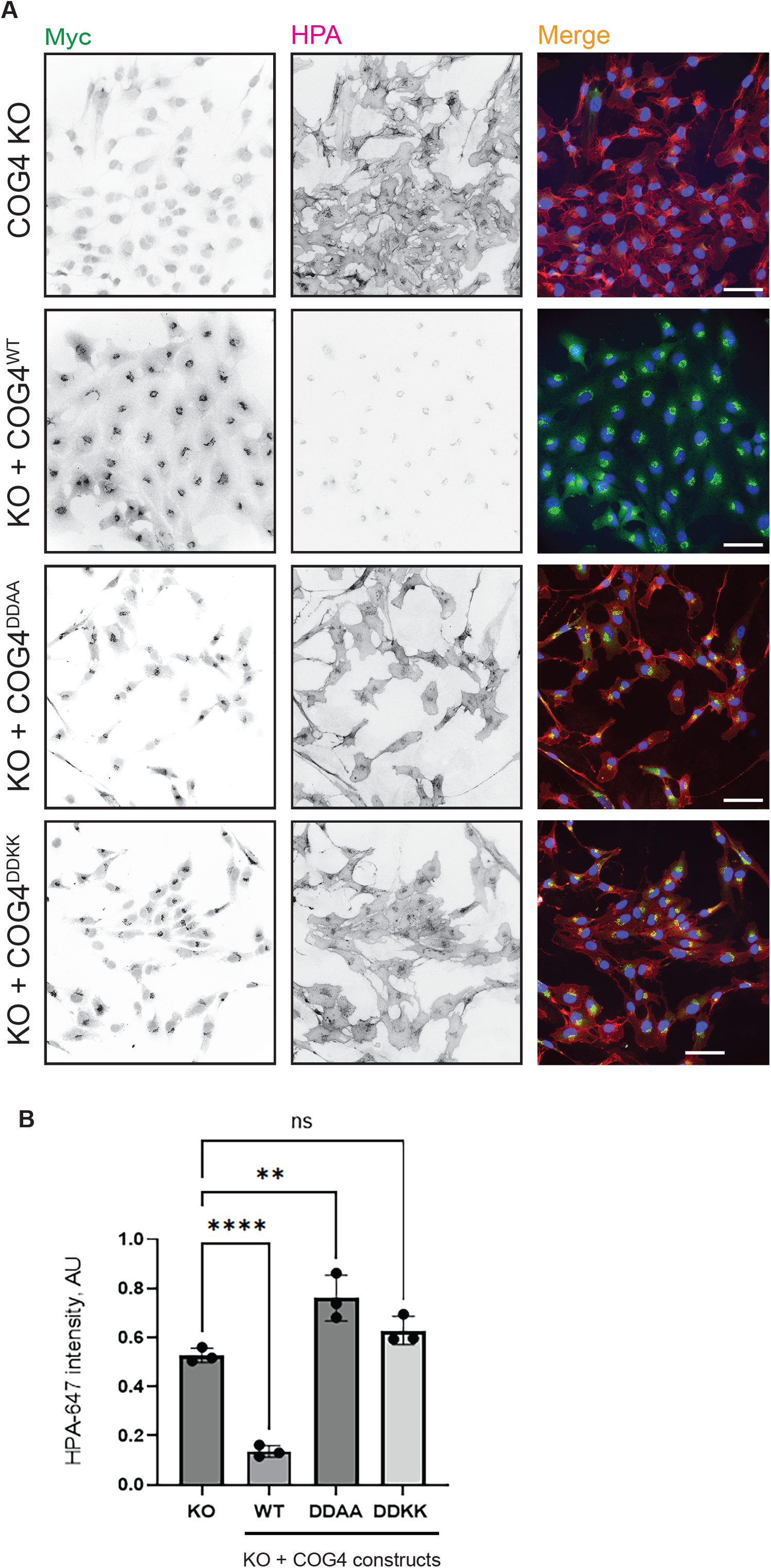
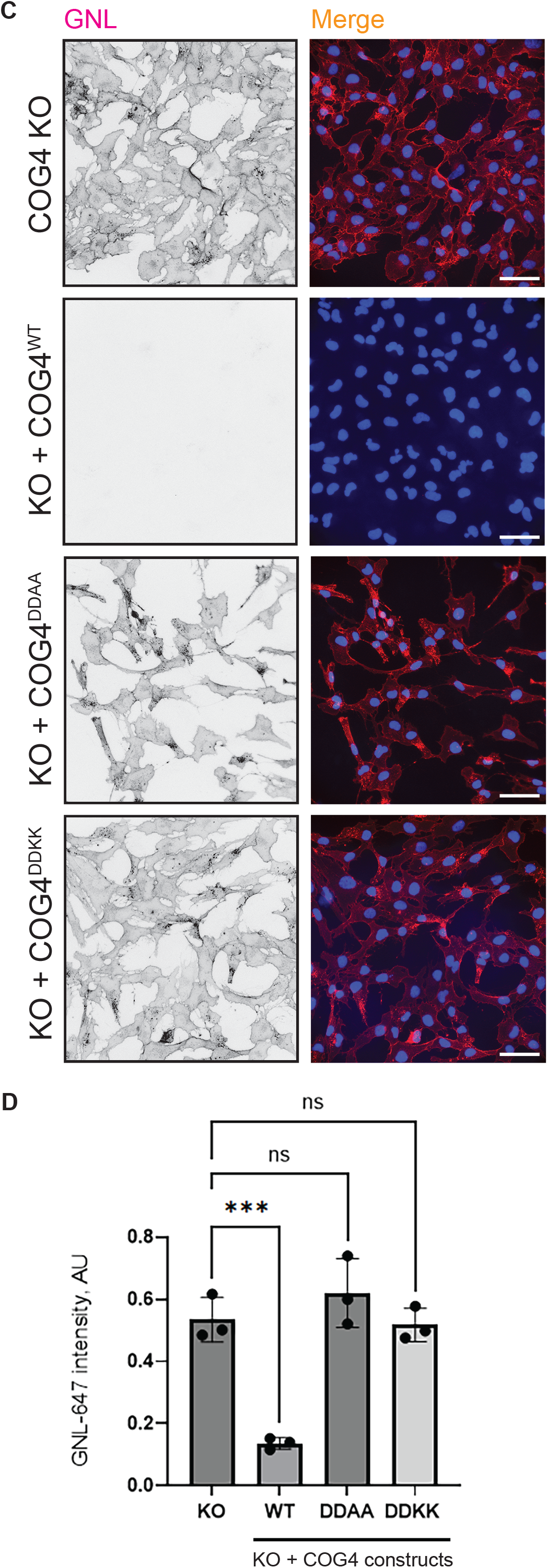
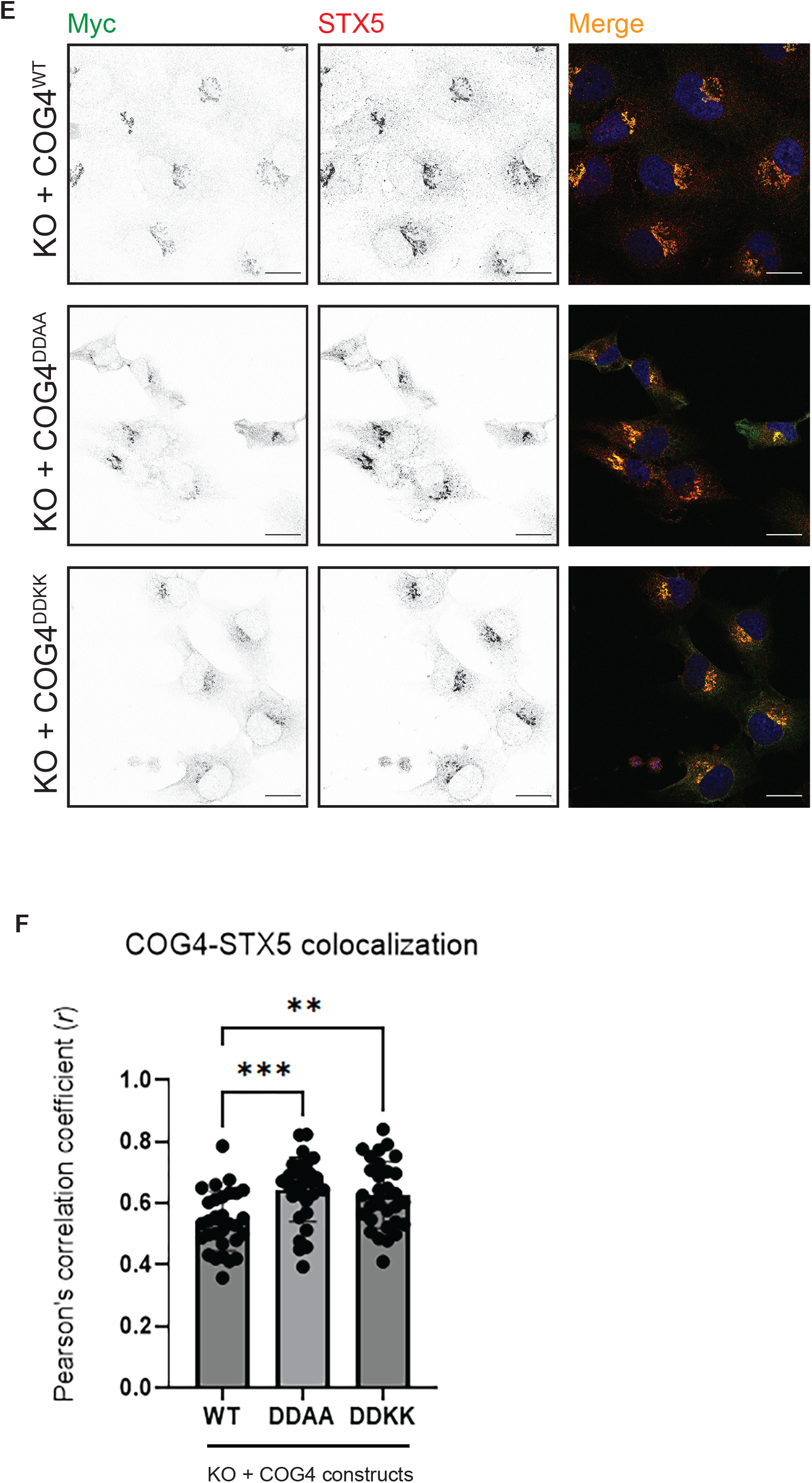
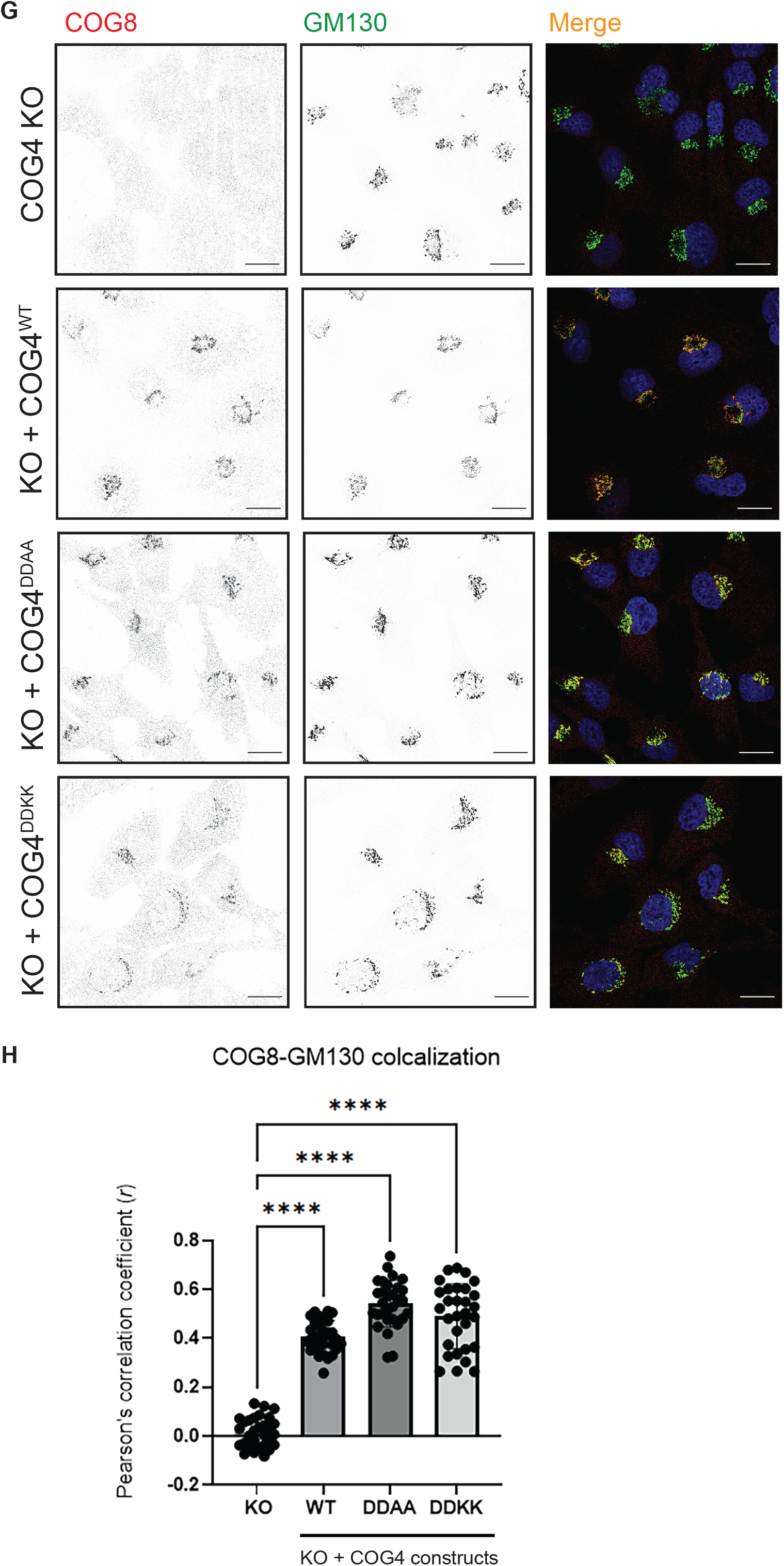
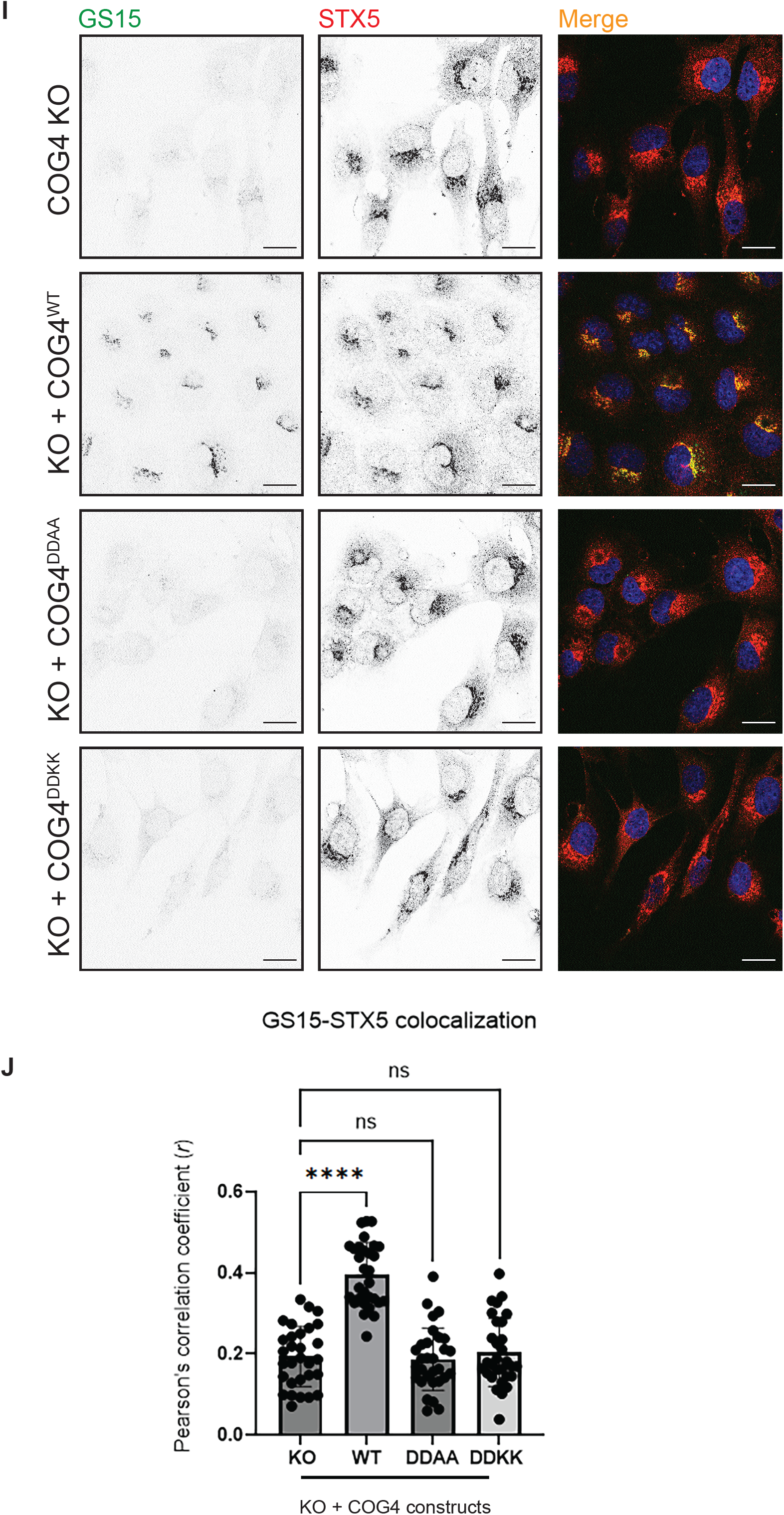
COG4 D452/D453 mutants retain Golgi localization but fail to restore normal glycosylation and GS15 localization. RPE1 COG4-knockout (KO) cells were complemented with myc-tagged wild-type COG4 (WT), COG4-D452A/D453A (COG4^DDAA^), or COG4-D452K/D453K (COG4^DDKK^), as indicated. Uncomplemented KO cells served as controls. (**A, C**) Representative fluorescence images of cells stained with HPA–647 (**A**) or GNL–647 (**C**) to assess alterations in O- and N-linked glycosylation, respectively. Anti-myc staining in panel A shows expression of the COG4 constructs throughout the complemented cell populations. (**B, D**) Quantification of HPA (**B**) and GNL (**D**) fluorescence. Mean fluorescence intensity was measured for each image using ImageJ and divided by the number of cells in that image. Values are expressed in arbitrary units (AU). (**E**) Confocal images of cells stained for myc and STX5, showing Golgi localization of WT and mutant COG4 proteins. (**F**) Quantification of myc–STX5 colocalization using Pearson’s correlation coefficient. (**G**) Confocal images of cells stained for COG8 and the Golgi marker GM130. (**H**) Quantification of COG8–GM130 colocalization using Pearson’s correlation coefficient. (**I**) Confocal images of cells stained for GS15/BET1L and STX5. STX5 retains its Golgi localization, whereas GS15 shows reduced Golgi localization in cells expressing the COG4 mutant. (**J**) Quantification of GS15–STX5 colocalization using Pearson’s correlation coefficient. Individual fluorescence channels are displayed in inverted grayscale; merged images show the indicated channels in color and nuclei in blue. Panels **A** and **C** were acquired using a 32× objective. Scale bar: 100 µm. Panels **E**, **G**, and **I** were acquired by confocal microscopy using a 63× objective. Scale bars: 20 µm. Bars show mean ± SD. Statistical analyses were performed using one-way ANOVA in GraphPad Prism. Comparisons are indicated by brackets. ns, not significant; **P < 0.01; ***P < 0.001; ****P < 0.0001.

To determine whether these mutations impair Golgi glycosylation, we stained the rescued cells with HPA and GNL. HPA staining at the cell surface and quantification across multiple fields showed that wild-type COG4 and COG4^2A^ reduced HPA binding to levels comparable to those of the wild-type rescue control. HPA binding remained elevated in cells expressing COG4^5A^ or COG4^7A^ and was comparable to that in uncomplemented KO cells (**Fig. 5A, B**). GNL staining showed the same pattern (**Fig. 5C, D**). Thus, the E277A/R281A substitutions preserved COG4’s ability to correct O- and N-glycosylation defects, whereas the five substitutions in the MUN-C-like subdomain prevented complementation despite preserved Golgi localization.

### COG4 interface mutants localize to the Golgi but fail to restore normal glycosylation

To determine the minimal amino acid residues required for the COG4-STX5 interaction, we focused on the conserved aspartates D452 and D453 in the MUN-C-like subdomain (**Fig. 2D, F**). These residues were also picked because other CATCHR proteins contain a conserved aspartate in their MUN-C-like subdomain (Pei et al., 2009). We replaced both residues with alanine to remove their negative charges (COG4^DDAA^) or with lysine to reverse their charges to introduce the steric clash (COG4^DDKK^). To enable future proximity labeling studies, we added TurboID to the C-terminus of each myc-tagged COG4 construct. We generated stable RPE1 COG4-KO cell lines expressing COG4^WT^–TurboID-myc (Aragon-Ramirez, Khakurel, Pokrovskaya, Sumya, & Lupashin, 2026), COG4^5A^–TurboID-myc, or the indicated mutants. (Aragon-Ramirez et al., 2026). HPA and GNL binding remained significantly elevated in cells expressing either mutant, both mutant rescues, relative to cells complemented with wild-type COG4, indicating that neither mutant corrected the O- or N-glycosylation defects (**Fig. 6A–D**). Anti-myc staining showed uniform expression of wild-type and mutant constructs across the respective cell populations (**Fig. 6A**).

Confocal microscopy showed that COG4^DDAA^ and COG4^DDKK^ retained Golgi localization, as assessed by colocalization of myc staining with the endogenous STX5 (**Fig. 6E, F**). Both mutants also restored Golgi localization of COG8, a lobe B subunit, as assessed by its colocalization with GM130 (**Fig. 6G, H**). The recovery of Golgi-localized COG8 suggests that both wild-type and mutant COG4 can assemble into a complex.

GS15/BET1L, a SNARE partner of STX5, showed significantly reduced colocalization with STX5 in cells expressing either mutant, even though STX5 retained its Golgi localization (**Fig. 6I, J**), a phenotype consistent with impaired assembly of STX5-containing SNARE complexes. Thus, mutating D452 and D453 is sufficient to prevent GS15 Golgi localization while maintaining proper COG complex assembly and localization. These results support that D452 and D453 in the MUN-C-like subdomain of COG4 are the minimal amino acids required for COG-dependent STX5 SNARE complex assembly.

### COG4^5A^ and COG4^DDAA^ show reduced coprecipitation with STX5

To determine whether substituting D452 and D453 within the predicted COG4–STX5 interface weakens their association, we transfected HEK293T cells stably expressing GFP–STX5 with myc-tagged wild-type COG4, COG4^5A^, or COG4^DDAA^ (D452A/D453A). We isolated GFP–STX5 and associated proteins using GBP beads. Both mutants showed significantly lower recovery with GFP–STX5 than wild-type COG4 (**Fig. 7A, B**), supporting a contribution of D452 and D453 to COG4–STX5 association. Consistent with these results, COG4^5A^–TurboID showed reduced proximity labeling of STX5 and GS15 while retaining labeling of COG3 (**Supp. Fig. 7**). These findings suggest that the interface mutations impair COG4 engagement with the Golgi SNARE machinery while preserving its proximity to another COG subunit.

**Figure 7.**
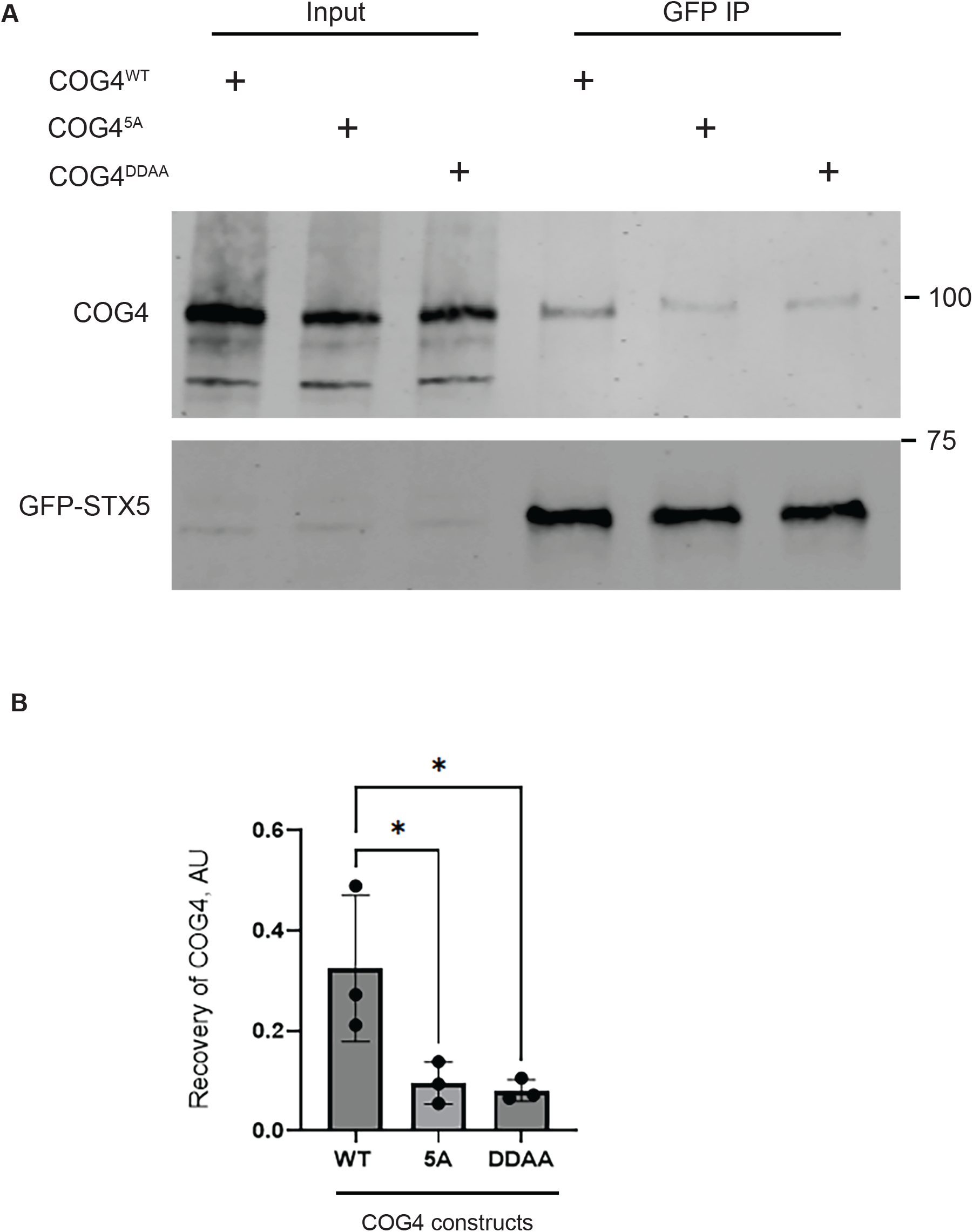
Mutations in the COG4 MUN-C-like subdomain reduce COG4 association with STX5. (**A**) WT HEK293T cells expressing GFP–STX5 were transiently transfected with plasmids encoding 3×myc-tagged COG4^WT^, COG4^5A^ (E442A/D452A/D453A/K459A/K460A), or COG4^DDAA^ (D452A/D453A). Following overnight expression, GFP–STX5 was isolated from cell lysates using GFP-binding protein beads. Input lysates and bound fractions (GFP IP) were analyzed by SDS–PAGE and immunoblotting with anti-myc and anti-GFP antibodies to detect COG4 constructs and GFP–STX5, respectively. Molecular masses are indicated in kDa. (**B**) Quantification of COG4 recovery with GFP–STX5. Anti-myc band intensities in the bound fractions were measured using LI-COR Image Studio and normalized to the corresponding anti-GFP signal. Values are expressed in arbitrary units (AU). Bars show the mean ± SD from three independent biological replicates, with individual replicates shown as points. Graphs were generated using GraphPad Prism 10. Statistical comparisons are indicated by brackets. *P < 0.05.

### Disruption of the COG4-STX5 interface causes Golgi structural defects resembling COG4 loss

Loss of COG4 has been shown to dramatically perturb Golgi morphology (Pokrovskaya et al., 2011). As shown in Figure 4A, mutations in the COG4-interacting region of STX5 result in comparable Golgi morphology disruption to that of the loss of COG4. To assess if D452 and D453 are directly involved in these observed phenotypes, we examined the Golgi ultrastructure in RPE1 COG4-KO cells stably expressing TurboID-tagged wild-type COG4, COG4^5A^, COG4^DDAA^, or COG4^DDKK^ by transmission electron microscopy. In COG4-KO cells, recognizable stacks were lost, and the Golgi region contained swollen cisternae and vacuole-like membrane profiles, resembling abnormalities previously reported in HEK cells lacking COG subunits (J. B. Blackburn, Pokrovskaya, Fisher, Ungar, & Lupashin, 2016), unlike the parental cells, which contain well-organized Golgi stacks. Wild-type COG4– TurboID restored stacked cisternae and near-normal Golgi morphology. In contrast, cells expressing COG4^5A^, COG4^DDAA^, or COG4^DDKK^ retained the knockout’s morphological defects (**Fig. 8A, B**).

**Figure 8.**
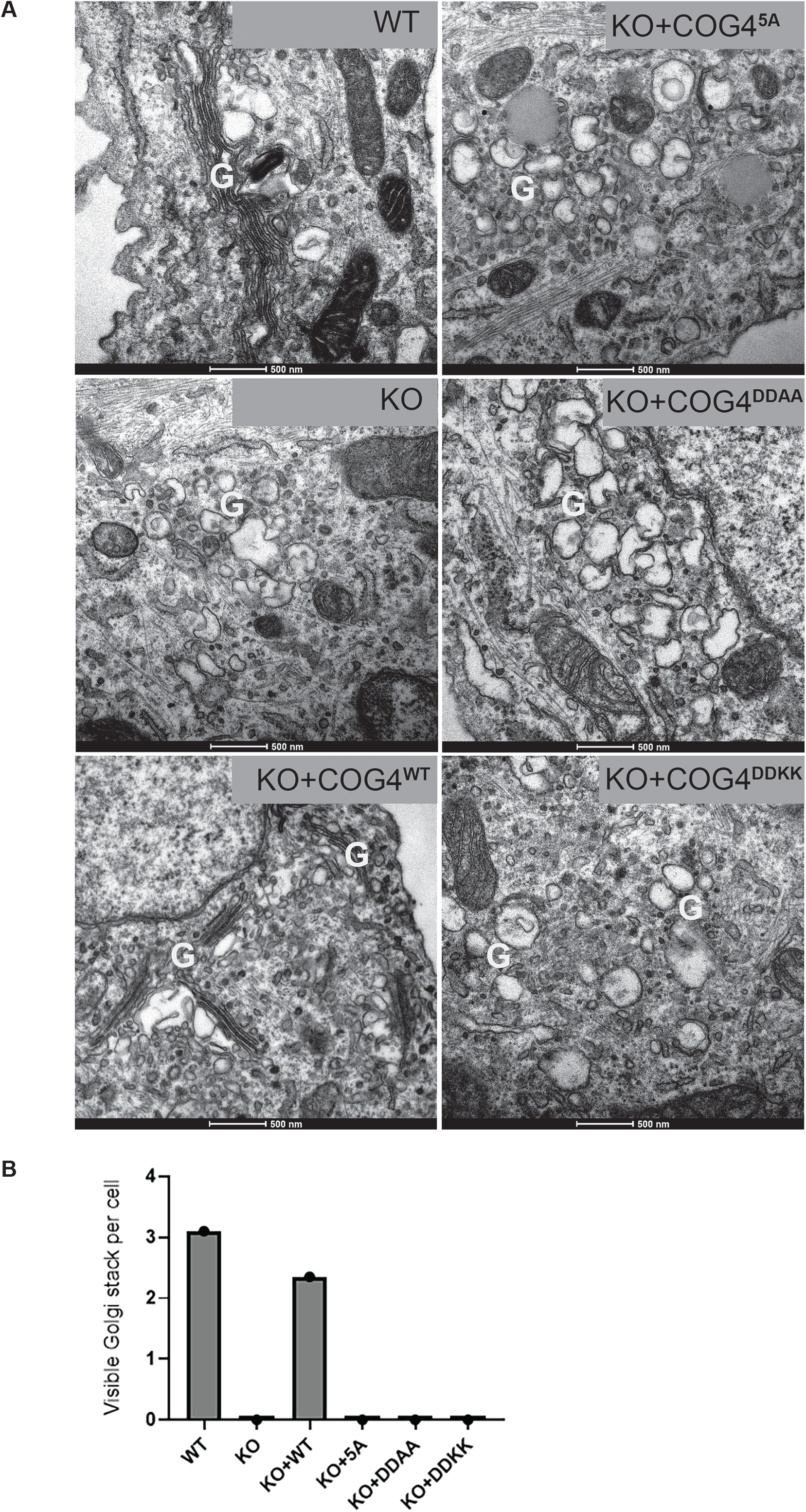
Mutations in the COG4 MUN-C-like subdomain prevent restoration of Golgi ultrastructure in COG4-knockout cells. **(A)** Representative TEM micrographs of wild-type RPE1 cells (WT), COG4-knockout cells (KO), and KO cells complemented with myc-tagged COG4^WT^ or the indicated mutants. COG4^5A^ contains E442A/D452A/D453A/K459A/K460A; COG4^DDAA^ contains D452A/D453A; and COG4^DDKK^ contains D452K/D453K. WT cells and KO cells complemented with COG4^WT^ exhibit stacked Golgi cisternae. In contrast, uncomplemented KO cells and cells expressing the COG4 mutants exhibit disrupted Golgi organization, with vesicular and dilated membrane profiles. G, Golgi region. Scale bars, 500 nm. **(B)** Quantification of visible Golgi stacks in the indicated cell populations.

Together, the complementation experiments identify D452 and D453 in the COG4 MUN-C-like subdomain as critical for Golgi function. Replacing these residues with either alanine or lysine preserved Golgi localization of COG4 and COG8 recruitment, but failed to restore normal glycosylation, GS15 localization, or Golgi cisternal stacking. The reduced recovery of COG4^DDAA^ with STX5 further links these functional defects to weakened COG4–STX5 association. These findings support a model in which the COG4–STX5 interface helps sustain retrograde trafficking within the Golgi and thereby maintain Golgi organization and glycosylation.

## Discussion

How vesicle tether proteins promote productive SNARE assembly remains a central question in membrane trafficking. Here, we identify a functionally important surface in the MUN-C-like subdomain of COG4 and implicate the Ha–Hb loop of STX5 in the same pathway. Earlier studies mapped associations with STX5 and its SM protein partner SCFD1/Sly1 to the N-terminal region of COG4 (Shestakova et al., 2007), (Laufman et al., 2009). Structural studies subsequently placed the COG4 N-terminus within the assembly core of lobe A, raising the question of how accessible the proposed binding sites are in the assembled complex (Laufman et al., 2009), (Ha et al., 2016). The N-terminal substitutions we tested did not reduce COG4 recovery with GFP–STX5 under overexpression conditions. In the same assay, the C-terminal fragment coprecipitated with STX5 more efficiently than full-length COG4. These results identify the C-terminal region as the major site for STX5 association and leave open how the two regions contribute to the intact complex.

Mutagenesis on both sides of the AF3-predicted interface linked these regions to Golgi function. Alanine or lysine mutation of the conserved COG4 residues D452 and D453 to alanine reduced COG4 recovery of STX5, while retaining Golgi localization of COG4 and recruitment of the lobe B subunit COG8. Additionally, this double mutation did not restore COG4 KO defects, such as impaired glycosylation, GS15 localization, or cisternal stacking. On the STX5 side, mutation of the Ha-Hb loop (3A) rescued GPP130 abundance and glycosylation more than mutation of the Habc-SNARE linker alone. The STX5^12M^ mutant (which incorporates both sets of mutations) retained Golgi localization and SCFD1 association but recovered less COG3 and GS15 and resulted in significant vesicle accumulation in the vicinity of the disorganized Golgi cisternae. Taken together, these data link the predicted interface to the assembly or maintenance of STX5-containing SNARE complexes and point to vesicle fusion as a likely step perturbed by the disruption.

Other CATCHR complexes engage SNAREs through structurally distinct contacts. In mammals, ZW10 and RINT1 are associated with the ER SNARE machinery containing STX18, BNIP1, p31/USE1, and SEC22B (Hirose et al., 2004). In yeast, the Dsl1 subunits Tip20 and Sec39 bind the N-terminal regulatory domains of the ER Qb-SNARE Sec20 and Qc-SNARE Use1, respectively (Travis et al., 2020), (Damico et al., 2024). Disrupting either tether–SNARE contact individually caused no detectable growth defect under the conditions tested. However, simultaneously disrupting a SNARE’s contact with its Dsl1 subunit and its contact with the other SNARE was lethal, indicating that the two contacts provide alternative routes for incorporating Sec20 or Use1 into the Dsl1–SNARE assembly (Damico et al., 2024). At the trans-Golgi network, mammalian VPS51 binds STX6, and its yeast counterpart binds Tlg1. Disrupting the Vps51–Tlg1 contact produced no detectable defect in the trafficking assays reported, leaving open whether other GARP–SNARE contacts compensate (Damico et al., 2024), (Fridmann-Sirkis, Kent, Lewis, Evans, & Pelham, 2006). Mammalian GARP depletion nevertheless reduced recovery of assembled STX16-containing SNARE complexes (Damico et al., 2024). In yeast, Sec6 binds assembled exocytic SNARE complexes, and reconstitution with purified octameric exocyst components stimulated several steps in SNARE assembly and membrane fusion (Dubuke, Maniatis, Shaffer, & Munson, 2015), (Lee et al., 2025). These studies establish several routes by which CATCHR complexes engage in fusion machinery. Whether they also share a conserved mode of Qa-SNARE recognition remains open.

Our findings prompt a reassessment of the previously proposed interactions of the COG4 N-terminal region with STX5 and SCFD1 (Shestakova et al., 2007), (Laufman et al., 2009). Multiple alanine substitutions within the proposed N-terminal STX5-binding region did not reduce COG4 coprecipitation with STX5 under the overexpression conditions tested (**Supp. Fig. 1**). Moreover, the C-terminal fragment containing the MUN-like region showed substantially greater recovery with GFP–STX5 than the N-terminal fragment (**Fig. 1**). Consistent with these biochemical results, AF3 models generated using proteins from six species placed the principal predicted STX5 interface within the COG4 MUN-like region (**Fig. 2** and data not shown).

Modeling the isolated COG4 N-terminal fragment provided further context for these observations. AF3 predictions of COG4 residues 1–153 with STX5 and SCFD1 yielded variable fragment placements and very low interface confidence (**Supp. Table 1**). Across five predictions, pairwise ipTM scores were 0.08–0.10 for COG4– STX5 and 0.12–0.14 for COG4–SCFD1, whereas the STX5–SCFD1 interface was consistently recovered with substantially greater confidence. Full-length COG4 models, by comparison, reproducibly positioned the MUN-C-like surface against the STX5 Ha–Hb loop and Habc–SNARE linker, encompassing the regions tested by mutagenesis. Combined with the functional defects caused by D452/D453 and Ha–Hb loop substitutions, these results favor the MUN-like region as the principal functional STX5-binding module of COG4. Additional N-terminal contacts may nevertheless depend on protein conformation or assembly context, particularly because the isolated fragment itself was modeled with low confidence.

**Table 1.** List of antibodies and lectins and their dilutions.

| Antibody | Source | Species | WB dilution | IF dilution |
|---|---|---|---|---|
| Myc tag | Cell signaling, #2276 | Mouse | N/A | 1:500 |
| Myc tag | Bethyl, #A190-205 A | Rabbit | 1:2000 | N/A |
| GFP tag | BioLegend, #902602 | Mouse | 1:2000 | N/A |
| STX5 |  | Rabbit | 1:500 | 1:500 |
| LAMP2 | DSHB, H4B4 | Mouse | 1:2000 | N/A |
| GPP130 | Covance, PRB-144C | Rabbit | 1:1000 | N/A |
| Actin |  | Mouse | 1:200 | N/A |
| HA tag | BioLegend, #901513 | Mouse | 1:2000 | N/A |
| COG3 | Lupashin lab | Mouse | 1:500 | N/A |
| COG8 | Sigma, SAB4200427 | Rabbit | N/A | 1:500 |
| GM130 | BD Biosciences, #610823 | Mouse | N/A | 1:500 |
| GM130 | CalBiochem, #CB1008 | Rabbit | N/A | 1:500 |
| GS15 | Lupashin lab | Rabbit | 1:500 | N/A |
| HPA lectin, Alexa Fluor 647 | Invitrogen, #L32454 | *Helix pomatia* | 1:2000 | 1:500 |
| GNL lectin, Alexa Fluor 647 | Lupashin lab |  | 1:2000 | 1:500 |
| Alexa Fluor 647 anti-Rabbit | Jackson Immuno Research/711-605-152 | Donkey | 1:4000 | N/A |
| Alexa Fluor 555 anti-Rabbit | Jackson Immuno Research/711-605-152 | Donkey | N/A | 1:1000 |
| Alexa Fluor 488 anti-Mouse | Jackson Immuno Research/715-545-151 | Donkey | N/A | 1:1000 |
| IRDye 800 anti-Mouse | LiCOR/926-68170 | Goat | 1:2000 | N/A |
| IRDye 800 anti-Rabbit | LiCOR/926-32211 | Donkey | 1:2000 | N/A |

AF3 modeling also placed a candidate SCFD1 contact within the COG4 MUN-D-like subdomain, suggesting an alternative to the reported SCFD1-binding site at COG4 residues 1–84 (**Fig. 2** and data not shown). We therefore tested the previously described E53A/E71A substitutions in our complementation system (Laufman et al., 2009). Two independent RPE1 COG4-KO clones expressing COG4-E53A/E71A–TurboID retained Golgi localization of the mutant protein and restored Golgi localization of GS15 and GS28/GOSR1 (**Supp. Fig. 8A–D**). Both clones also corrected the elevated HPA and GNL binding of uncomplemented knockout cells, with rescue comparable to that provided by wild-type COG4–TurboID (**Supp. Fig. 8E–G**). The restoration of Golgi SNARE localization differs from the phenotype previously reported in HeLa cells (Laufman et al., 2009). Differences in cellular background, expression level, or experimental design could account for this discrepancy, including the use of acute siRNA-mediated depletion in the earlier study versus stable knockout and complementation here. Adaptation during stable cell-line selection may also contribute. Our results establish that E53 and E71 are dispensable for the Golgi functions measured under these conditions. Direct biochemical analysis of SCFD1 binding, together with comparisons in matched cellular systems, will be needed to define the contributions of the proposed N-terminal and MUN-D-like interfaces.

The structural similarity between CATCHR subunits and the Munc13 MUN domain suggests a testable model for how COG4 could prime STX5. In neuronal fusion, Munc13 promotes the transition of Munc18-bound syntaxin 1 from its closed state toward SNARE assembly. This activity depends on the conserved MUN-B NF pocket and is sensitive to substitutions in the syntaxin 1 linker (Li et al., 2011), (Yang et al., 2015), (Magdziarek et al., 2020), (Wang et al., 2020). We did not identify an equivalent NF pocket in the CATCHR structures examined. The COG4–STX5–SCFD1 model instead accommodates COG4 binding to a closed-like STX5 conformation and predicts a local rearrangement of the Habc–SNARE linker. COG4 could therefore engage SCFD1-bound STX5 before SNARE assembly and position the linker for a subsequent transition. This model would explain how COG4 can influence SNARE assembly while remaining compatible with SCFD1 association. Measuring STX5 conformation and SNARE assembly kinetics in a purified system would distinguish an effect on syntaxin opening from effects on partner recruitment or complex stability.

The comparative AF3 models suggest that the MUN-like regions of VPS53 and RINT1 may also be contacting the regulatory domains of their respective Qa-SNAREs. The complementation and proximity labeling data we’ve seen so far (data not shown) sufficiently support testing these predicted interfaces directly.

In the separate model of the STX5 SNARE complex with SCFD1 and COG4, the COG4 MUN-D-like subdomain is positioned close to the N-terminal regulatory domain of the Qb-SNARE GOSR1. That arrangement supports the idea that COG4 could use adjacent subdomains to position both STX5 and GOSR1 for assembly. Future studies will test these contacts individually to determine whether COG4 acts as a primary primer of STX5, organizes a Q-SNARE intermediate, or acts at both stages.

## Methods Cell culture

HEK293T (WT or COG4 KO) and hTERT-RPE1 cells (WT or COG4 KO) were grown on DMEM/F12 (1:1) (1X) (Dulbecco’s Modified Eagle Medium, Gibco) supplemented with 10% fetal bovine serum (FBS) (Gibco, Cat# 16000-044). Cells were incubated at 37 °C in a 5% CO2, 90% humidity-controlled incubator. For passage, cells were trypsinized with 0.25% Trypsin-EDTA (1X) (Gibco) at room temperature for 2 minutes and resuspended in DMEM-F12 medium supplemented with 10% FBS.

### AlphaFold3 and ChimeraX

We obtained amino acid sequences from UniProt (Bateman et al., 2024) and submitted them to the AlphaFold3 server (Abramson et al., 2024) to predict structures of individual proteins and protein complexes. We selected models with the highest confidence scores reported by AlphaFold3 and imported them into UCSF ChimeraX for visualization and figure preparation (Meng et al., 2023) . Models were rendered as ribbons or molecular surfaces and oriented to highlight structural features and residues at the predicted protein–protein interfaces

### Plasmid preparation

GenScript synthesized and sequence-verified all STX5 and COG4 mutant constructs. Human STX5 isoform 2 was tagged at its N-terminus with GFP and co-expressed with HA-tagged SCFD1 from a single open reading frame using a p2A sequence. To confer siRNA resistance, we replaced the STX5 target sequence 5′-TAGCCTCAACAAACAAATT-3′ with 5′-CTCCCTCAATAAACAAATT-3′. The resulting SCFD1–HA–P2A–GFP–STX5 cassette was subcloned into pENTR1A and then transferred by recombination into a pLenti vector containing the STX5 promoter (D’Souza et al., 2023). Wild-type and mutant human COG4 constructs carried either a C-terminal 3×myc tag or a C-terminal TurboID–3×myc tag. The constructs were subcloned into pENTR1A and then transferred by recombination into a pLenti vector containing the COG4 promoter (Sumya et al., 2021). The pLenti plasmids were sequenced by Plasmidsaurus for verification. DNA constructs are listed in Table 1.

### Stable cell line productions

Stable cell lines were generated by lentiviral transduction as previously described (Sakuma, Barry, & Ikeda, 2012), (Khakurel, Kudlyk, & Lupashin, 2023). To produce lentivirus, cells were grown on collagen-coated plates to 50–60% confluence and co-transfected with the pLenti construct and three helper plasmids (pMD2.G, pRSV-Rev, and pMDLg-pRRE) (Cribbs, Kennedy, Gregory, & Brennan, 2013; Sakuma et al., 2012), using Lipofectamine in DMEM supplemented with FBS. After 16 h, the medium was replaced with 8 mL of Opti-MEM containing 1× GlutaMAX and 1 mM sodium butyrate. The following day, the first collection of virus-containing medium was harvested, centrifuged at 400 × g for 10 min, and passed through a 0.45-μm polyethersulfone (PES) filter. The filtered supernatant was then added to recipient cells grown in six-well plates: HEK293 and RPE1 COG4-knockout cells for the COG4 constructs, and wild-type HEK293 cells for the STX5 constructs. Three days after transduction, cells were transferred to 10-cm dishes for antibiotic selection. HEK293 cells expressing STX5 constructs were selected with 100 μg/mL hygromycin. Cells expressing COG4 constructs were selected with G418 at 400 μg/mL for HEK293 cells or 500 μg/mL for RPE1 cells. Following selection, cells were maintained in 20 μg/mL hygromycin for STX5-expressing lines or 100 μg/mL G418 for COG4-expressing lines. Expression of homogeneity was assessed by IF. Transduced RPE1 cells showed homogeneous COG4 expression and did not require sorting. HEK293 populations showed heterogeneous expression and were therefore sorted. Cells expressing GFP-tagged STX5 were sorted by GFP fluorescence using flow cytometry. HEK293 COG4-KO cells expressing mutant COG4-Myc were single-cell sorted into 96-well plates, and clones were screened by WB and IF to select the final cell lines.

### Coimmunoprecipitation assay

HEK293 cells stably co-expressing SCFD1–HA and GFP–STX5 or expressing GFP–TurboID (GFP–TID) as a specificity control, were seeded in 12-well plates. At 50–60% confluence, cells were transfected with plasmids encoding full-length COG4 or its N-terminal or C-terminal fragments, each fused to a C-terminal 3×myc tag. The following day, cells were lysed on ice for 20 min in 400 µL of lysis buffer containing 1% Triton X-100, 20 mM HEPES (pH 7.5), 100 mM NaCl, 1 mM dithiothreitol, 5 mM MgCl₂, 1 mM PMSF, and 1× Halt protease inhibitor. Lysates were clarified by centrifugation at 13,000 × g for 10 min at 4°C. A 50-µL aliquot of the supernatant was reserved as input, and 330 µL was incubated with GFP-binding protein (GBP) beads (7.5 µL packed bead volume) for 90 min at room temperature with rotation (Willett, Pokrovskaya, Kudlyk, & Lupashin, 2014). Beads were pelleted at 1,000 × g for 1 min, and the supernatant was collected as the flow-through. Beads were washed twice with lysis buffer lacking PMSF and the protease inhibitor Halt, followed by two washes with 1× PBS. During the first PBS wash, beads were transferred to a fresh tube. Bound proteins were eluted in 30 µL of 2× Laemmli sample buffer containing 7% β-mercaptoethanol by heating at 95°C for 5 min. Input and flow-through samples were mixed with 6× Laemmli sample buffer containing 7% β-mercaptoethanol and heated under the same conditions. Input, flow-through, and bound fractions were analyzed by Western blotting. For each bound fraction, the COG4 signal was normalized to the corresponding GFP-tagged bait signal to account for differences in bait recovery. Where indicated, normalized values were expressed as fold changes relative to the wild-type control. Coimmunoprecipitation assays were performed in triplicate, and data were plotted using GraphPad Prism 10.

### Western blotting

Proteins were separated on 4–15% gradient Mini-PROTEAN TGX gels (Bio-Rad) and transferred to 0.2-μm nitrocellulose membranes (Amersham Protran, Cytiva) using a Pierce G2 Fast Blotter (Thermo Scientific). Membranes were blocked for 30 min in EveryBlot blocking buffer (Bio-Rad, catalog no. 12010020), then incubated with primary antibodies diluted in the same buffer for 1 h at room temperature or overnight at 4°C. After four 4-min washes in PBS, membranes were incubated for 1 h with fluorescent secondary antibodies diluted in a 1:1 mixture of blocking buffer and PBS. Primary and secondary antibodies are listed in Table 2. Membranes were washed four more times for 4 min each in PBS, then stained with Ponceau S for 2 min to assess total protein loading. For HPA and GNL lectin blotting, membranes were blocked with 3% BSA in TBS for 30 min or with LI-COR blocking buffer, then incubated with HPA-647 or GNL-647 for 30 min. Blots were imaged using an Odyssey Imaging System, and band intensities were quantified by densitometry using LI-COR Image Studio software.

**Table 2.** DNA constructs used in this study.

| Plasmid | Source | Reference |
|---|---|---|
| pLenti COG4<sup>WT</sup> -3xmyc | Lupashin lab | This study |
| pLenti COG4<sup>3A</sup> -3xmyc | Lupashin lab | This study |
| pLenti COG4<sup>4A</sup> -3xmyc | Lupashin lab | This study |
| pLenti COG4<sup>7A-1</sup> -3xmyc | Lupashin lab | This study |
| pLenti COG4<sup>7A-2</sup>-3xmyc | Lupashin lab | This study |
| pLenti COG4<sup>2A</sup> -3xmyc | Lupashin lab | This study |
| pLenti COG4<sup>5A</sup> -3xmyc | Lupashin lab | This study |
| pLenti COG4<sup>7A</sup> -3xmyc | Lupashin lab | This study |
| pLenti COG4<sup>5A</sup> -TurboID-3xmyc | Lupashin lab | This study |
| pLenti COG4<sup>DDAA</sup> -TurboID-3xmyc | Lupashin lab | This study |
| pLenti COG4<sup>DDKK</sup> -TurboID-3xmyc | Lupashin lab | This study |
| pLenti SCFD1-HA p2A GFP-STX5<sup>WT</sup> | Lupashin lab | This study |
| pLenti SCFD1-HA p2A GFP-STX5<sup>3A</sup> | Lupashin lab | This study |
| pLenti SCFD1-HA p2A GFP-STX5<sup>8GS</sup> | Lupashin lab | This study |
| pLenti SCFD1-HA p2A GFP-STX5<sup>9GS</sup> | Lupashin lab | This study |
| pLenti SCFD1-HA p2A GFP-STX5<sup>12M</sup> | Lupashin lab | This study |
| pEntra no ccDB (w48-1) | Addgene #17398 | Gift from Eric Campeau & Paul Kaufman |
| pEntra hCOG4-TID Myc | Lupashin lab | (Aragon-Ramirez et al., 2026) |
| pEntra hCOG4 Myc | Lupashin lab | (Sumya et al., 2021) |
| pMD2.G | Addgene #12259 | Gift from Didier Trono |
| pMDLg/pRRE | Addgene #12251 | Gift from Didier Trono |
| pRSV-Rev | Addgene #12253 | Gift from Didier Trono |
| pLenti STX5 long pr DEST Hygro | Lupashin lab | (D'Souza et al., 2023) |
| pLenti COG4 long pr NEO DEST | Lupashin lab | (Sumya et al., 2021) |

### siRNA knockdown

Endogenous STX5 was depleted using a previously published STX5-targeting siRNA (TAGCCTCAACAAACAAATT; (D’Souza et al., 2023)) at a final concentration of 50 nM. Wild-type HEK293 cells expressing siRNA-resistant GFP–STX5 mutants 2A, 8GS, or 12M, together with wild-type HEK293 cells as a control, were grown in 12-well plates to 10–20% confluence. On day 1, cells were transfected with the siRNA using Lipofectamine RNAiMAX (Invitrogen) in Opti-MEM and incubated overnight. The following day, the medium was replaced with DMEM/F-12 supplemented with 10% FBS. A second transfection with the same siRNA at a final concentration of 50 nM was performed on day 3. On day 5, cells were lysed in hot 2% SDS. Lysates were sonicated using a Sonifier Cell Disrupter (model W185D) and heated at 95°C for 5 min. Before electrophoresis, samples were mixed with 2× sample buffers, heated at 95°C for 10 min, and analyzed by Western blotting.

### Immunofluorescence analysis

Cells were plated on glass coverslips to 80%–90% confluency and fixed with 4% PFA diluted in PBS for 15 min at room temperature. Cells were then permeabilized with 0.1% Triton X-100 in PBS for 1 min, followed by a 50 mM ammonium chloride wash for 5 min to remove remaining fixative. Then, cells were blocked with 1% BSA and 0.1% saponin in PBS for 20 min for subsequent incubation with primary antibody (diluted in 1% cold fish gelatin, 0.1% saponin in PBS) for 45 min, washed with PBS, and then incubated with fluorescently conjugated secondary antibodies for 30 min and thoroughly washed with PBS to remove unbound antibody. Coverslips were then mounted on glass microscope slides using Prolong Gold antifade reagent (Life Technologies). Cells were imaged with a 63× oil 1.4 numerical aperture objective on an LSM880 Zeiss Laser inverted microscope and an Airyscan super-resolution confocal microscope using ZEN software. Quantitative analysis of single-slice confocal images was performed using ImageJ or ZENBlue version 2.6. All the microscopic images shown are Z-stacked Maximum Intensity Projection images.

### Transmission electron microscopy (TEM)

Samples were processed for TEM according to Valdivia’s lab protocol (Cocchiaro & Valdivia, 2009) with some modifications (Pokrovskaya et al., 2012). Briefly, the excess growth medium was removed from the dish with cells, and an equal volume of 1x fixative was added for 5 min incubation. Then the 0.5x fixative was replaced with 1x fixative for 10 min at RT. The 1x fixative was composed of 4% PFA (EMS) and 1% GA (EMS) in 0.1 M Phosphate buffer, pH 7.4. Finally, cells were fixed on ice for 20 min in 2.5% glutaraldehyde and 0.05% malachite green (EMS) prepared in 0.1 M sodium cacodylate buffer (pH 6.8). Samples were then post-fixed for 30 min at room temperature in 0.5% osmium tetroxide and 0.8% potassium ferricyanide in 0.1 M sodium cacodylate buffer, followed by incubation in 1% tannic acid for 20 min on ice and 1% aqueous uranyl acetate for 1 h at room temperature. Specimens were dehydrated through a graded ethanol series, washed with Propylene Oxide (EMS), and incubated in 50% PO/ resin mixture before embedding in Araldite 502/Embed 812 resins (EMS). Ultrathin sections (50 nm) were prepared using a Leica UltraCut UCT ultramicrotome and post-stained with aqueous uranyl acetate and Reynolds’ lead citrate (EMS). Images were acquired on an FEI Tecnai TF20 transmission electron microscope operated at 80 kV and equipped with an FEI Eagle 4k digital camera controlled by FEI acquisition software. Vesicles were counted per photo taken in Golgi areas at 14500x. The graph was made using GraphPad Prism 10.

### Statistical analysis

All results are representative of at least three biological replicates. For statistical analysis of lectin intensity, one-way ANOVA was performed in GraphPad Prism 10. The remaining graphs were also made using GraphPad Prism 10. Error bars represent standard deviation.

## Acknowledgements

We thank our colleagues who generously provided reagents used in this study. We are particularly grateful to the members of the Lupashin laboratory for valuable discussions and critical feedback. We also thank the UAMS Digital Microscopy for access to their facilities and for their expert technical support.

## Funding

This work was supported by the NIH/NIGMS grant R01GM083144 for Vladimir Lupashin.

---

Supplementary Figure 1. Analysis of candidate STX5-binding residues in the COG4 N-terminal region. **(A)** Two views of the AlphaFold2-predicted structure of human COG4 residues 1–222, shown as a surface representation with charged residues labeled. **(B)** Alanine substitution mutants targeting the indicated charged residues, along with surface representations of the 7A-1 and 7A-2 variants. **(C)** GFP–STX5 pulldown analysis of wild-type COG4 (WT) and the indicated N-terminal mutants co-expressed in HEK293T cells. Input lysates and GFP-bound fractions (GFP IP) were analyzed by SDS–PAGE and immunoblotting for COG4 and GFP–STX5. Molecular masses are indicated in kDa. **(D)** Quantification of COG4 recovery normalized to GFP–STX5 recovery in the bound fractions. Values are expressed in arbitrary units (AU), which represent the ratio between the mean signal intensity of COG4 and GFP-STX5. Individual measurements are shown as points.

Supplementary Figure 2. S**t**ructural **similarity between the UNC13A MUN domain and COG4. (A, B)** AlphaFold-predicted structures of UNC13A residues 920–1394 **(A)** and COG4 residues 134–785 **(B)**, visualized as ribbon representations using UCSF ChimeraX. MUN subdomains A–D are indicated. Superposition of Cα traces within the aligned regions of UNC13A (blue) and COG4 (red). Structural alignment was performed using the TM-align server, yielding a root-mean-square deviation (RMSD) of 4.91 Å and a TM-score of 0.67077. Since UNC13A fragment is shorter, the calculation is normalized to the length of the shorter fragment to prevent the larger COG4 fragment from artificially penalizing.

Supplementary Figure 3. S**t**ructural **model of the COG4–STX5 interface. (A)** Overview of the AlphaFold3-predicted COG4–STX5 complex, illustrated relative to a schematic Golgi membrane. COG4 MUN-like subdomains A–D are shown as surface representations. STX5 isoform 2 is shown as a ribbon, with its N-terminal region, Habc domain, linker, SNARE motif, and C-terminal membrane anchor indicated. The domain schematic below the model shows the corresponding residue ranges. **(B)** Enlarged view of the proposed interface between COG4 MUN-C and STX5, highlighting COG4 residues D452, D453, K459, and K460, the STX5 Ha–Hb loop (residues 66–74), and linker residues R153, R154, and E155. Dashed lines illustrate proposed contacts. STX5 residue numbering corresponds to isoform 2.

Supplementary Figure 4. S**T**X5 **mutants retain Golgi localization. (A)** Representative fluorescence images of HEK293 cells stably expressing GFP-tagged human STX5 isoform 2, either wild-type (WT) or the indicated mutants (3A, 8GS, 9GS, and 12M). Cells were stained for the cis-Golgi marker GM130. Individual channels are shown in inverted grayscale. Merged images show GFP–STX5 in green, GM130 in red, and nuclei in blue. **(B)** GFP–STX5 and GM130 colocalization were quantified in triplicate using Pearson’s correlation coefficient (*r*). None of the mutants differed significantly from WT in the indicated comparisons (ns, not significant).

Supplementary Figure 5. R**e**covery **of SCFD1, COG3, and GS15 with wild-type and mutant STX5 in the presence or absence of N-ethylmaleimide.** WT HEK293T cells stably expressing GFP–STX5, either wild-type (WT) or the 12M mutant, were left untreated or treated with 1 mM N-ethylmaleimide (+NEM) for 10 min at 37 °C before lysis. The STX5^12M^ mutant combines replacement of Ha–Hb loop residues 66–74 with a Gly/Ser sequence and the R153A/R154A/E155A substitutions. GFP–STX5 was isolated from cell lysates using GFP-binding protein beads. Input lysates and bound fractions (GFP IP) were analyzed by SDS–PAGE and immunoblotting to detect GFP–STX5, HA-tagged SCFD1, endogenous COG3, and GS15/BET1L. The asterisk indicates a nonspecific band. Molecular masses are indicated in kDa. Blots are representative of three independent biological replicates.

Supplementary Figure 6. G**o**lgi **localization of COG4 interface mutants. (A)** Representative fluorescence images of RPE1 COG4-knockout cells complemented with myc-tagged wild-type COG4 (WT) or the indicated mutants (2A, 5A, and 7A), stained for myc and the Golgi marker GM130. Mutant definitions are provided in Figure 2. Individual channels are displayed in inverted grayscale. Merged images show myc in red, GM130 in green, and nuclei in blue. Scale bars, 10 µm. **(B)** Quantification of COG4–GM130 colocalization using Pearson’s correlation coefficient. Bars show mean ± SD, with individual measurements shown as points. Statistical analysis was performed using ANOVA. Comparisons with WT are indicated by brackets. ns, not significant; ***P < 0.001;****P < 0.0001.

Supplementary Figure 7. P**r**oximity **labeling of COG and SNARE machinery by wild-type and COG4^5A^– TurboID.** RPE1 cells stably expressing GFP–TurboID (negative control) and RPE1 COG4-KO cells stably expressing wild-type COG4–TurboID or COG4^5A^–TurboID were subjected to biotin proximity labeling for 15 min. The COG4-5A mutant contains E442A/D452A/D453A/K459A/K460A substitutions in the MUN-C-like subdomain. Biotinylated proteins were enriched from cell lysates using streptavidin beads. Input lysates and streptavidin-bound fractions were analyzed by SDS–PAGE and immunoblotting with antibodies against TurboID, COG3, STX5, GS15/BET1L, and SCFD1. STX5 L and STX5 S indicate the endogenous long and short isoforms, respectively. Molecular masses are indicated in kDa.

Supplementary Figure 8. T**h**e **E53A/E71A substitutions preserve COG4-mediated rescue of Golgi SNARE localization and glycosylation.** Experiments were performed in hTERT-RPE1 COG4-KO cells (CTRL) or the same cells stably expressing myc-tagged wild-type COG4–TurboID (COG4 WT) or the COG4-E53A/E71A– TurboID mutant. Two independent mutant clones were analyzed (E53/71A Cl1 and Cl2). **(A)** Immunoblot analysis of COG4–TurboID expression using anti-myc antibodies. Actin serves as a loading control. **(B–D)** Representative immunofluorescence images of cells stained for the cis-Golgi marker GM130 (red) together with myc-tagged COG4 **(B)**, GS15/BET1L **(C)**, or GS28/GOSR1 **(D)** in green. Both mutant clones retain Golgi localization of COG4 and restore the Golgi localization of GS15 and GS28. **(E)** Representative images of cells stained with HPA–647 or GNL–647, shown in orange, with nuclei in blue. Wild-type COG4 and both mutant clones suppress the elevated lectin binding observed in uncomplemented knockout cells. Scale bars, 10 µm. **(F, G)** Lectin blots probed with fluorescently labeled HPA–647 **(F)** or GNL–647 **(G)** to assess alterations in O- and N-linked glycosylation, respectively. Signals were detected by fluorescence scanning.

## Abbreviations

CATCHR: Complexes Associated with Tethering Containing Helical Rods
CCT: Coiled-Coil Tether
COG: Conserved oligomeric Golgi complex
GARP: Golgi-Associated Retrograde Protein
EARP: Endosome-Associated Recycling Protein
IF: Immunofluorescence
KD: Knockdown
KO: Knockout
MS: Mass Spectrometry
NRZ: NAG, RINT1, and ZW10
PM: Plasma Membrane
SM: Sec1/Munc18
SNARE: Soluble N-ethylmaleimide-sensitive factor Attachment protein Receptor
STX5: Syntaxin 5
WB: Western blot

## Notes

### Competing Interest Statement

The authors have declared no competing interest.

