## Supplementary Figures 1-8 for "A Conserved Interface Between COG4 and Syntaxin 5 Is Essential for Intra-Golgi Membrane Trafficking"

Supplementary Figure 1

**A**

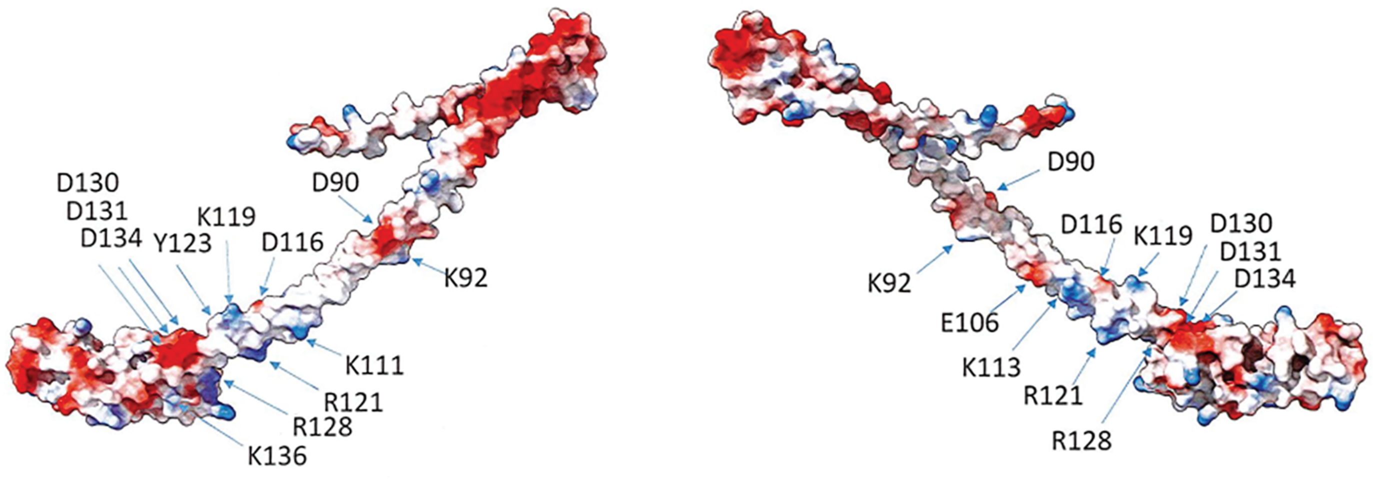


**B**

COG4$^{3A}$ = D116A K119A D130A
COG4$^{4A}$ = E106A K111A R113A R121A
COG4$^{7A-1}$ = D90A K92A E106A K111A R113A D116A K119A
COG4$^{7A-2}$ = R121A Y123A R128A D130A D131A D13A K136A

**C**

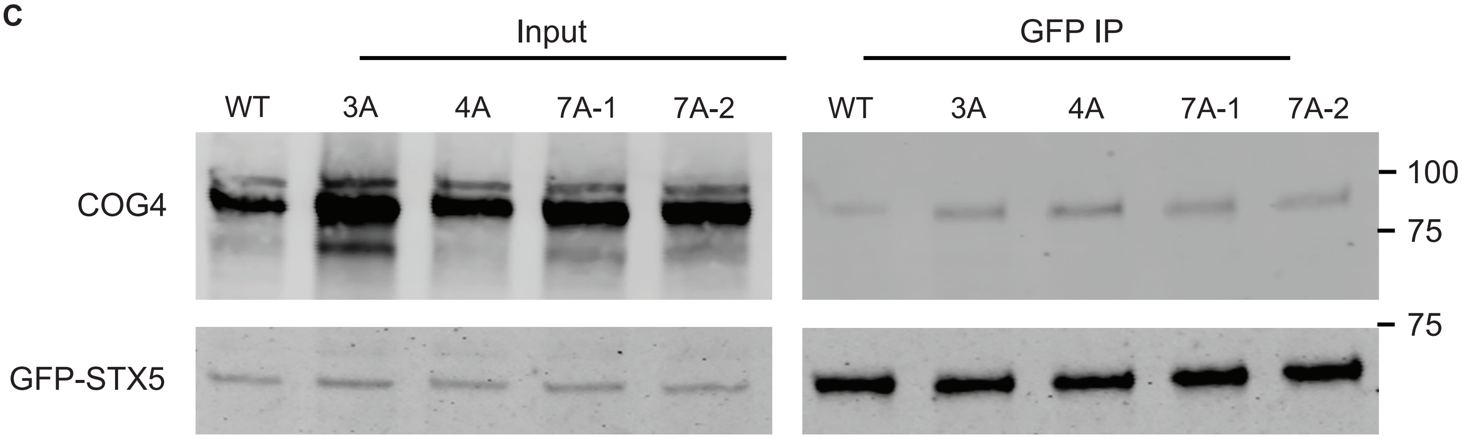


**D**

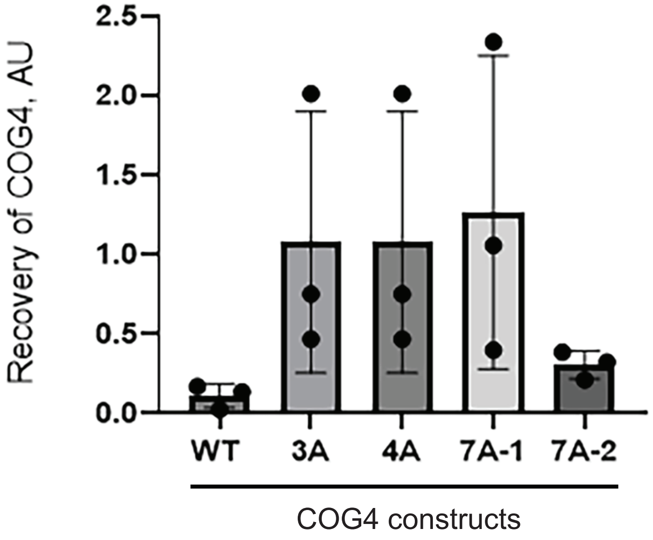

Supplementary Figure 2

**A**

**UNC13A 920-1394**

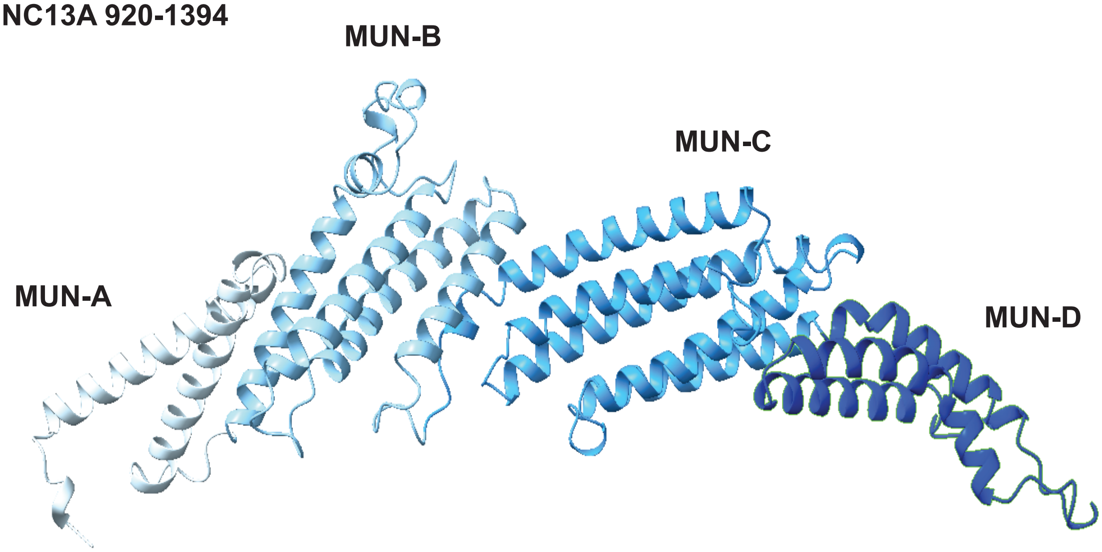


**B**

**COG4 134-785**

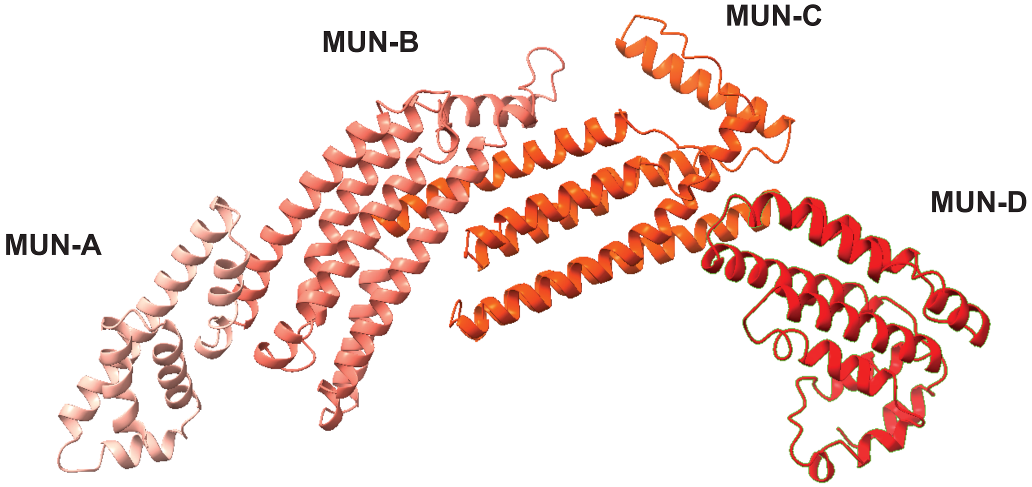


**C**

**Superposed carbon traces in the aligned regions**

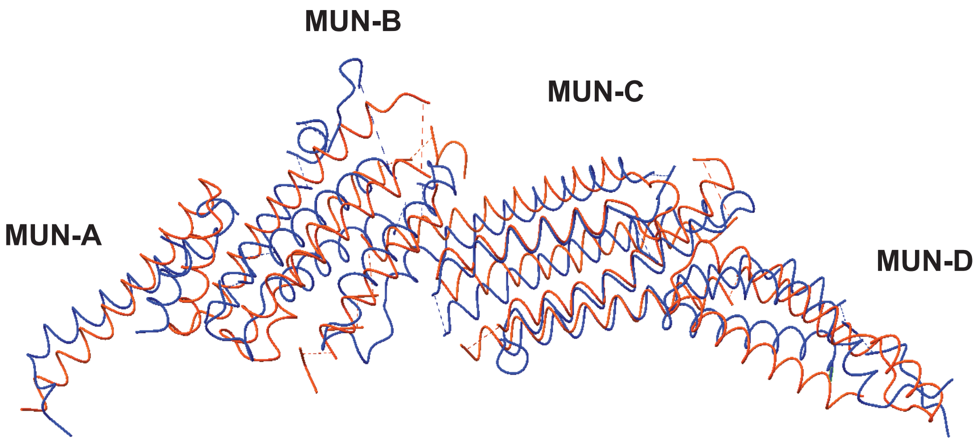

Supplementary Figure 3

A

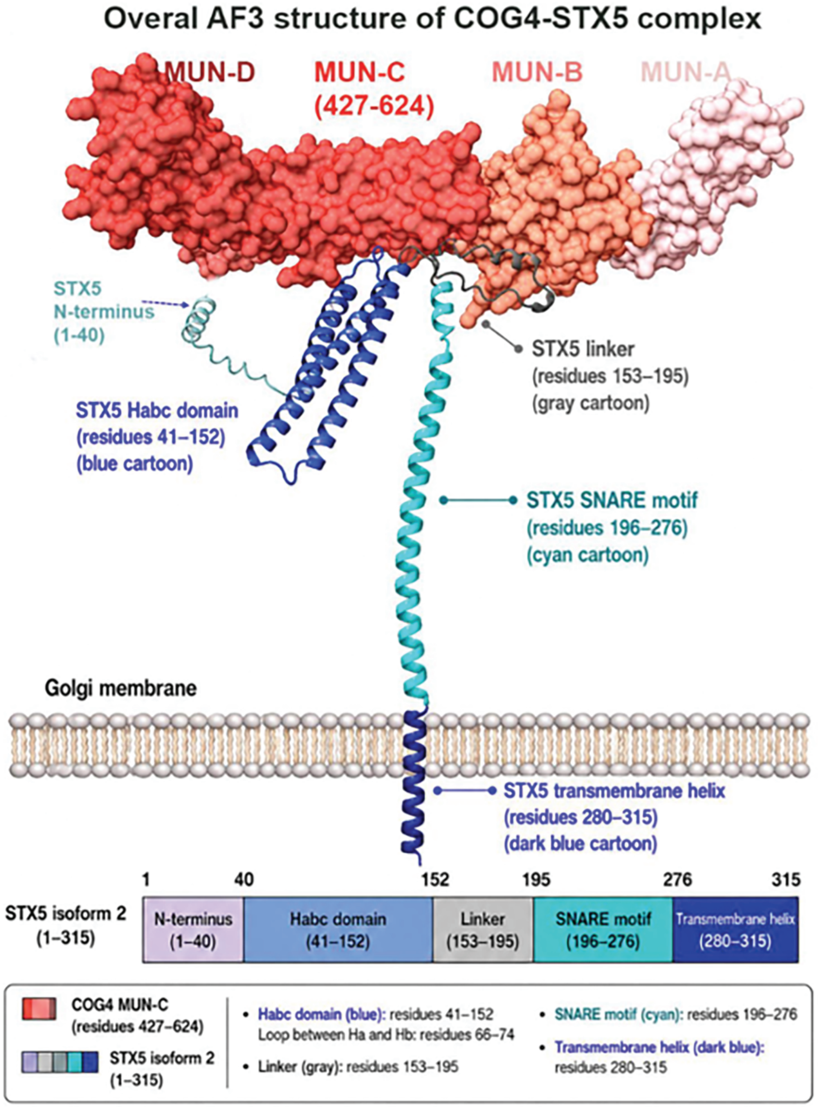


B

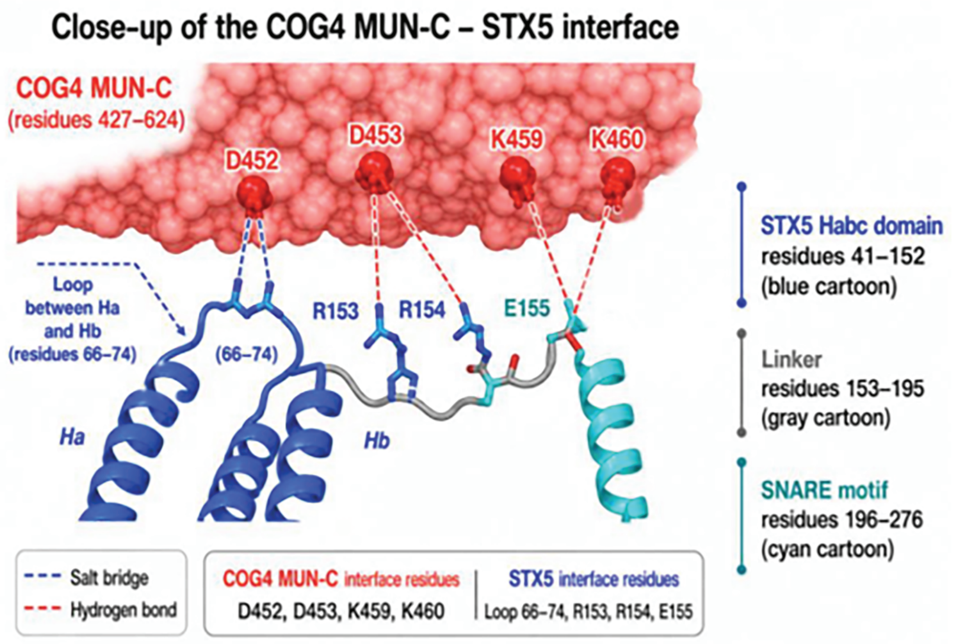

Supplementary figure 4

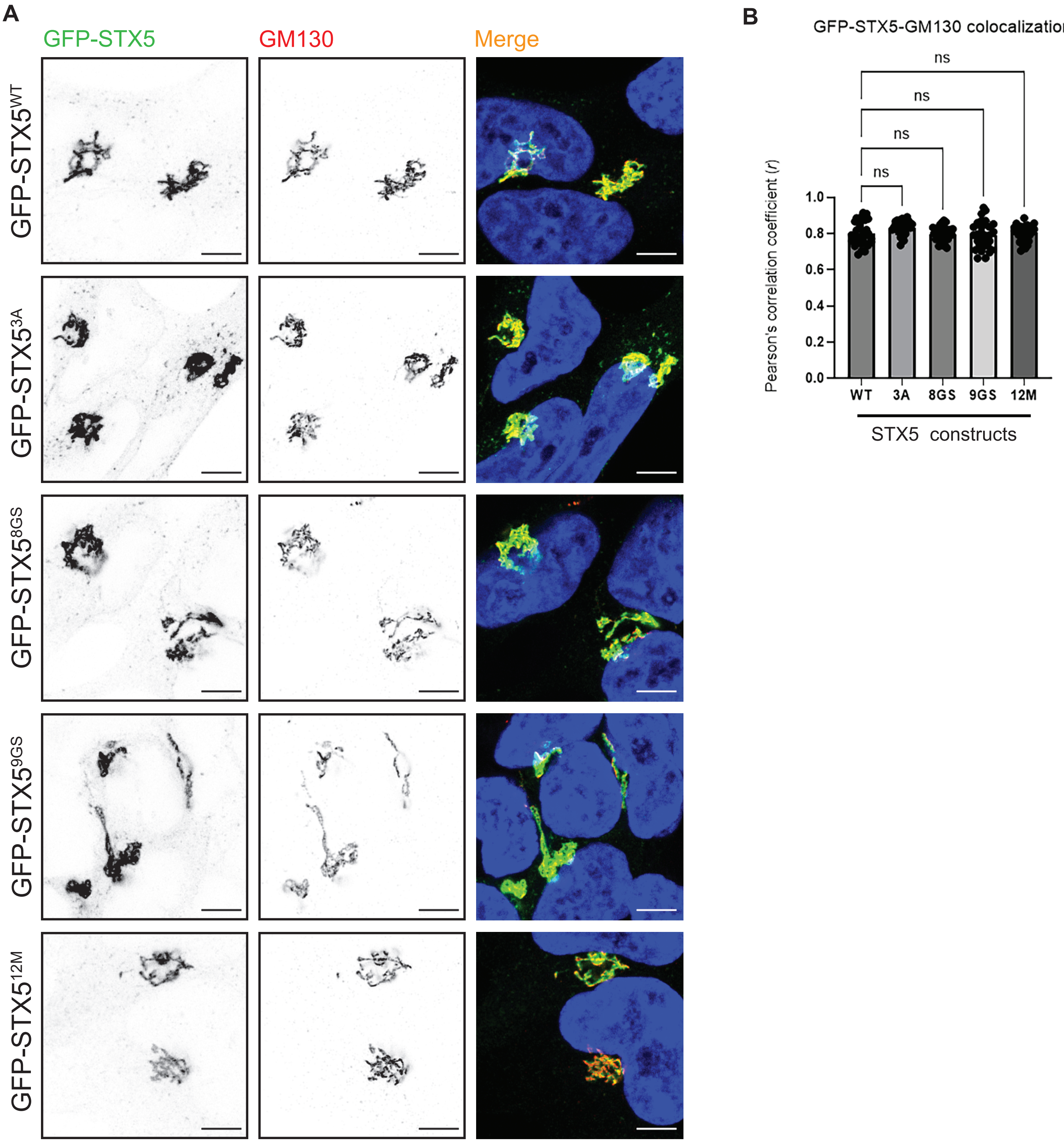

Supplementary Figure 5

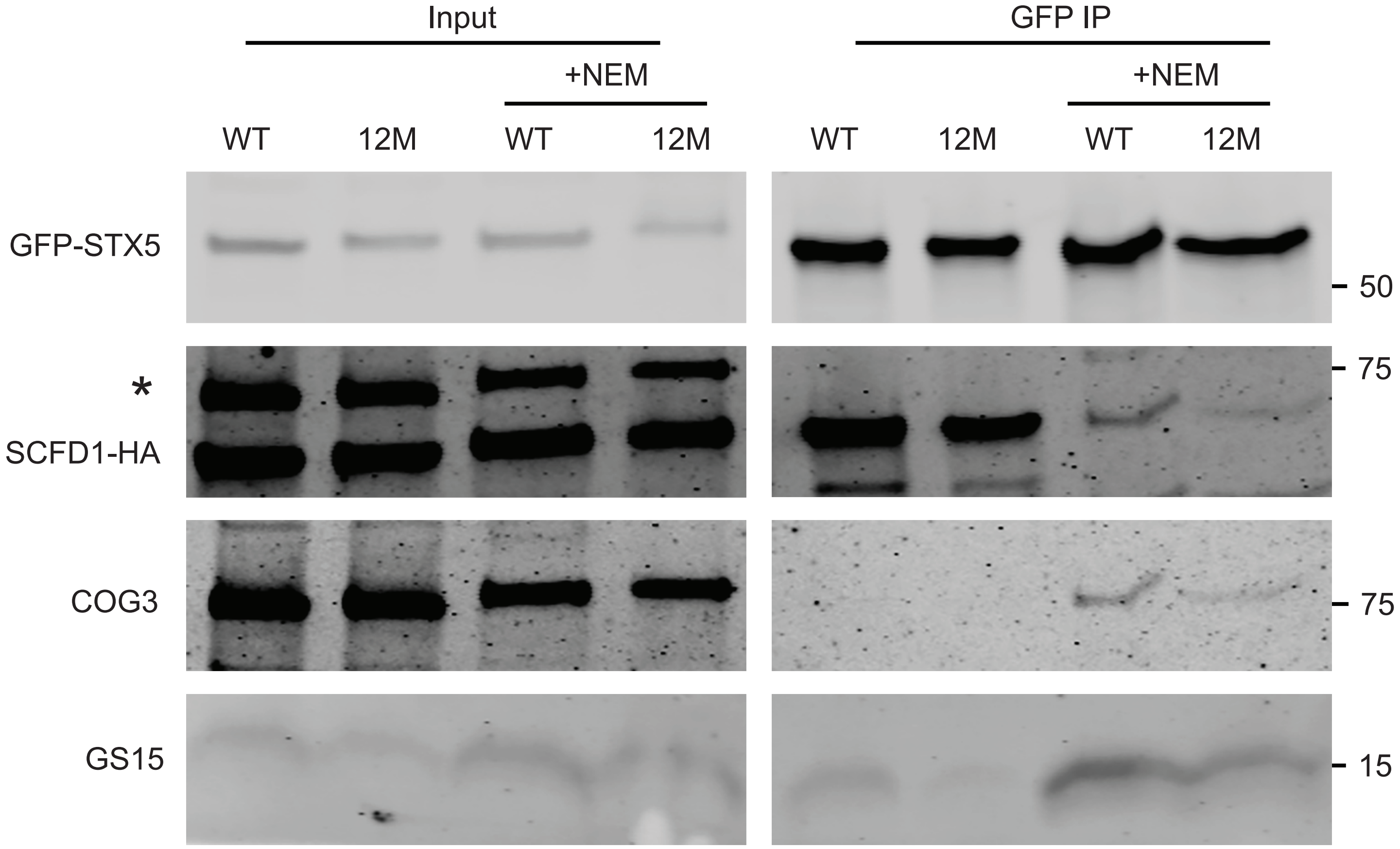

Supplementary Figure 6

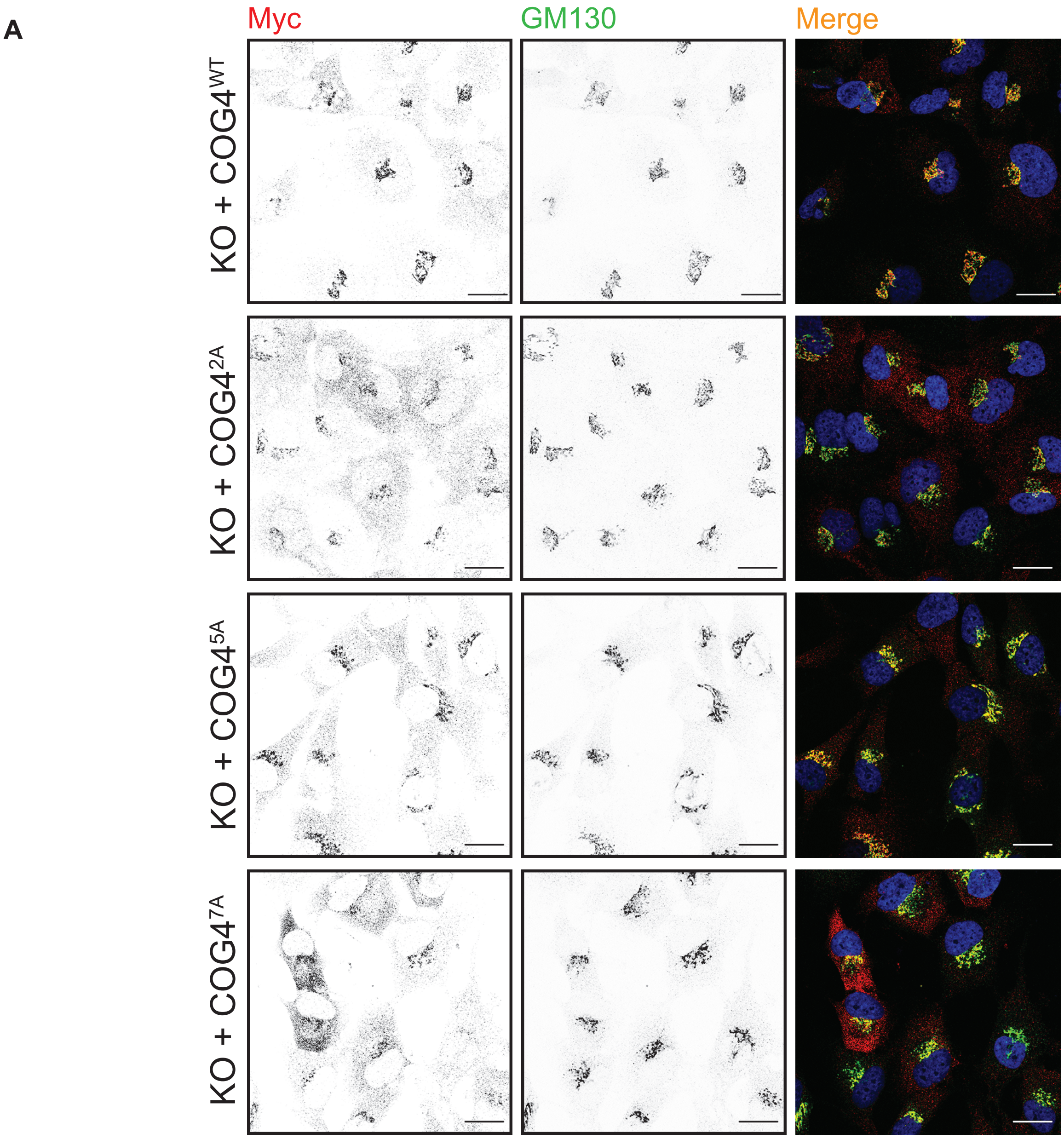


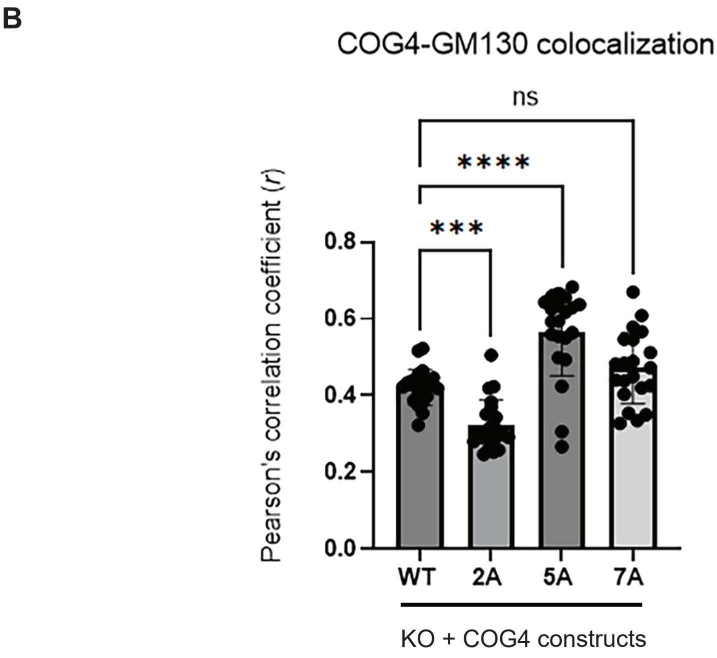

Supplementary Figure 7

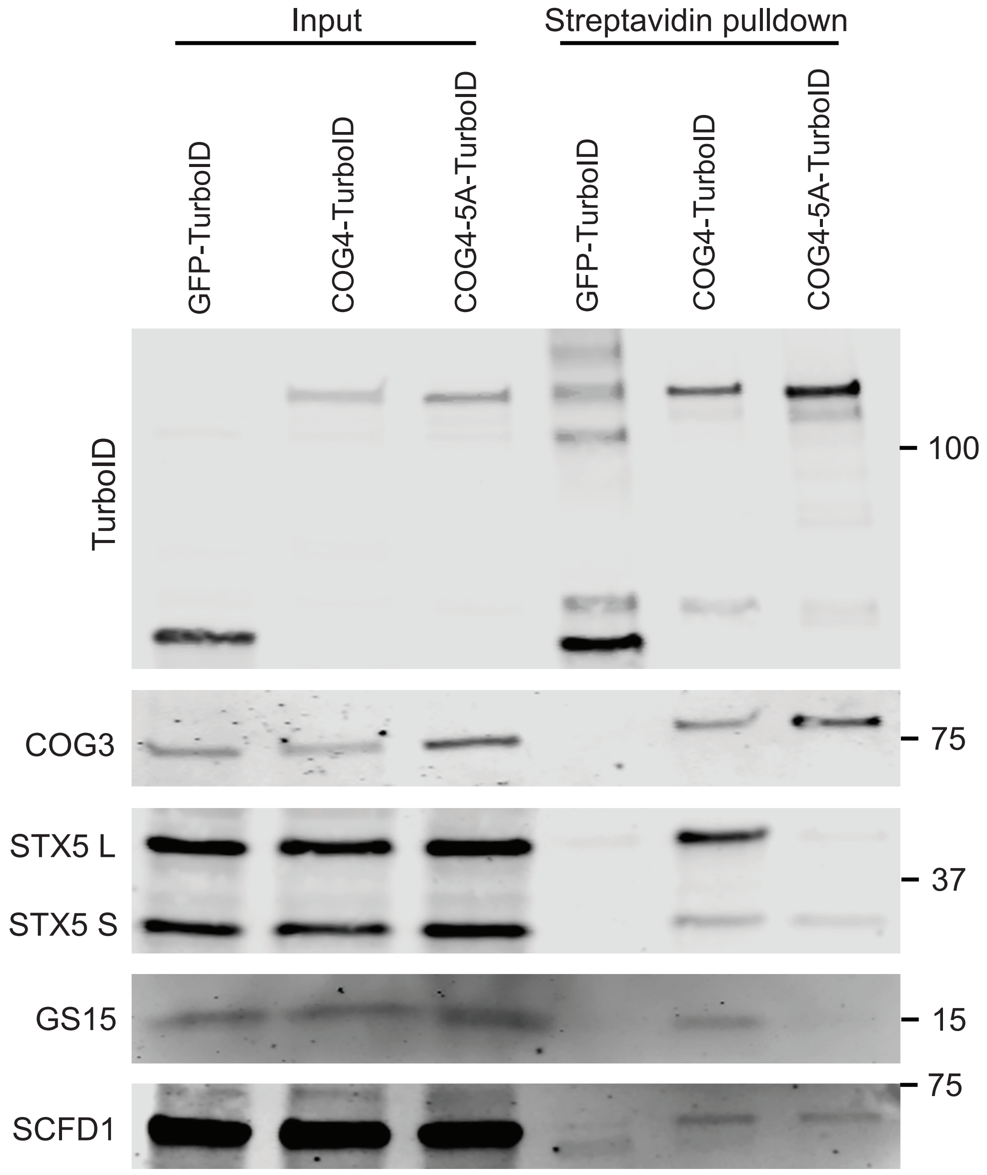

Supplementary Figure 8

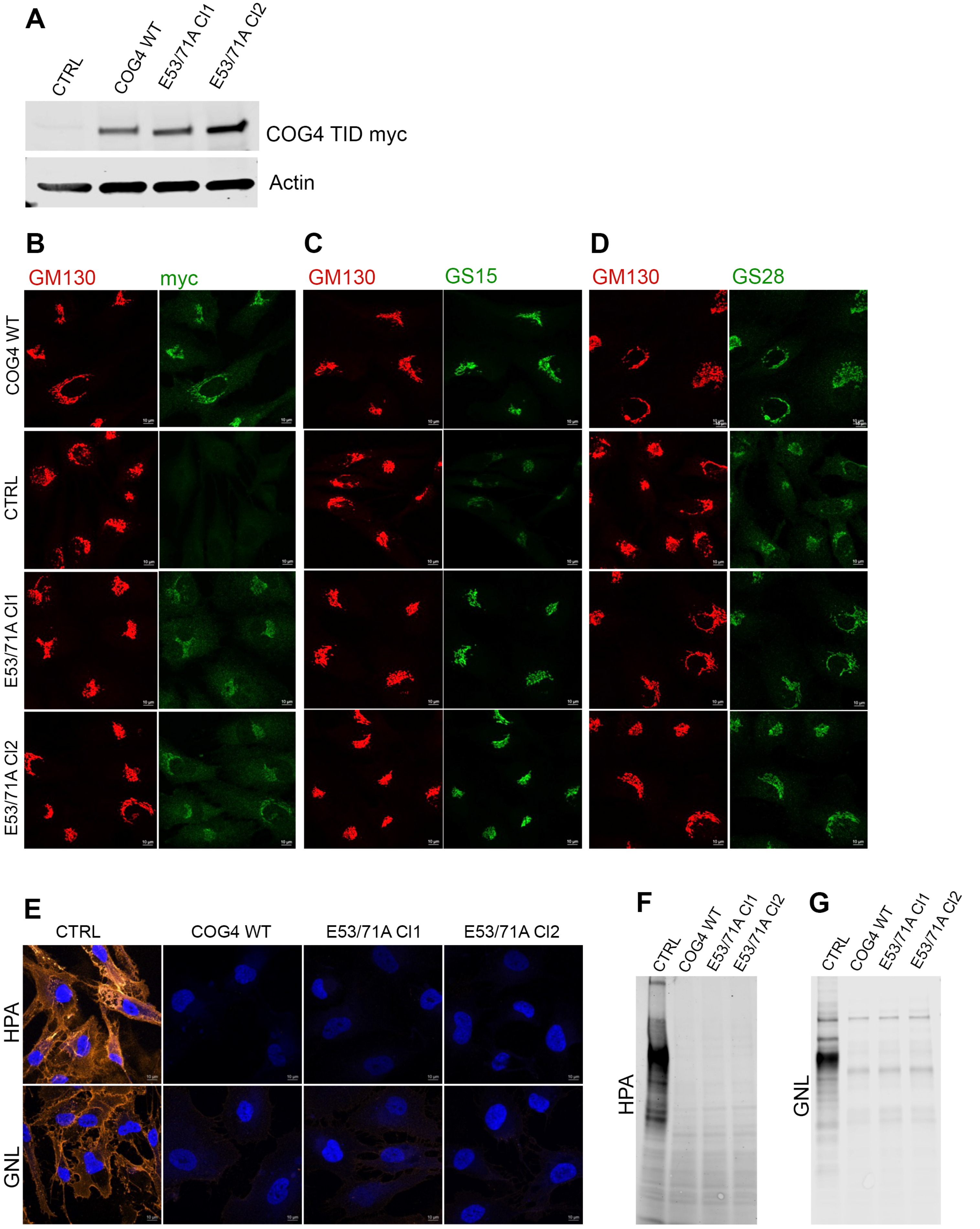
