## Supplementary Table 1 for "A Conserved Interface Between COG4 and Syntaxin 5 Is Essential for Intra-Golgi Membrane Trafficking"

**Supplementary Table 1. AlphaFold3 confidence metrics for complexes containing N-terminal or full-length COG4**

| Confidence metric | COG4(1–153)–STX5–SCFD1 | COG4(1–785)–STX5–SCFD1 |
|---|---|---|
| COG4–STX5 pairwise ipTM | 0.08–0.10 | 0.56–0.57 |
| COG4–SCFD1 pairwise ipTM | 0.12–0.14 | 0.55–0.57 |
| STX5–SCFD1 pairwise ipTM | 0.66–0.67 | 0.68–0.70 |
| Minimum interchain PAE, COG4→STX5 (Å) | 20.5–23.4 | 2.3–2.4 |
| Minimum interchain PAE, COG4→SCFD1 (Å) | 21.3–23.3 | 6.4–6.6 |
| Mean COG4 Cα pLDDT | 49.8–52.9 | 78.8–79.4 |

AlphaFold3 models were generated using either the COG4 N-terminal fragment (residues 1–153) or full-length COG4 (residues 1–785), together with identical STX5 and SCFD1 sequences. Values represent the minimum–maximum range across five models generated for each complex. Pairwise interface predicted template modeling scores (ipTM, scale 0–1) describe confidence in the relative placement of each chain pair, with higher values indicating greater confidence. Predicted aligned error (PAE) describes uncertainty in relative residue placement, with lower values indicating greater confidence. Minimum interchain PAE was calculated separately for each model from the PAE matrix block containing COG4 rows and STX5 or SCFD1 columns, as indicated by the arrows. Mean COG4 predicted local distance difference test scores (pLDDT, scale 0–100) were calculated across Cα atoms, with higher values indicating greater local structural confidence. Mean pLDDT values cover the complete COG4 sequence included in each prediction.
